# Mapping ovarian cancer spatial organization uncovers immune evasion drivers at the genetic, cellular, and tissue level

**DOI:** 10.1101/2023.10.16.562592

**Authors:** Christine Yiwen Yeh, Karmen Aguirre, Olivia Laveroni, Subin Kim, Aihui Wang, Brooke Liang, Xiaoming Zhang, Lucy M. Han, Raeline Valbuena, Sylvia Katina Plevritis, Michael C. Bassik, Michael P. Snyder, Brooke E. Howitt, Livnat Jerby

## Abstract

Immune exclusion and evasion are central barriers to the success of immunotherapies and cell therapies in solid tumors. Here we applied single cell spatial and perturbational transcriptomics alongside clinical, histological, and genomic profiling to elucidate immune exclusion and evasion in high-grade serous tubo-ovarian cancer (HGSC). Using high-plex spatial transcriptomics we profiled more than 1.3 million cells from 95 tumors and 60 patients, revealing generalizable principles in HGSC tumor tissue organization. Our data demonstrates that effector T cells resist stroma-mediated trapping and sequestration. However, upon infiltration into the tumor, T cells, as well as Natural Killer (NK) cells, preferentially co-localize only with a subset of malignant cells that manifest a distinct transcriptional cell state. The latter consists of dozens of co-regulated genes and is repressed under various copy number alterations. Performing CRISPR Perturb-seq screens in ovarian cancer cells, we identified functionally diverse genetic perturbations – including knockout of the insulin sensing repressor *PTPN1* and the epigenetic regulator *ACTR8* – that de-repress the proposed immunogenic malignant cell state identified in patients and indeed sensitize ovarian cancer cells to T cell and NK cell cytotoxicity. Taken together, our study uncovered a profound connection between somatic genetic aberrations, malignant cell transcriptional dysregulation, and immune evasion at the cellular and tissue level, allowing us to identify targets that reprogram malignant cell states as an avenue to unleash anti-tumor immune responses.

## INTRODUCTION

Multicellular dysregulation plays a key role in the initiation and progression of a wide range of diseases, including cancer, where tumor development and accompanying immune responses depend on (and shape) the location of diverse cell type populations, tissue properties, and organization (1–7). Cellular and animal models have been instrumental in recovering central immune suppressors (8–10) and led to major breakthroughs in cancer patient care. However, many cancer patients do not respond to current immunotherapies (11–13), resulting, at least in part, from two central gaps. First, in contrast to the study of cancer genetics, where genome sequencing of tumors across large and diverse patient populations provided a strong foundation to study the genetic basis of cancer and develop targeted therapies, we still lack equivalent maps of tumor tissue organization to study the inherently spatial processes of multicellular dynamics and immune exclusion in patients. Second, identifying the regulators of cell states and reciprocal intercellular interactions poses additional challenges and requires functional interrogation across a larger search space of combinatorial gene-environment perturbations.

In tubo-ovarian high grade serous carcinoma (HGSC) – the most common and aggressive form of ovarian cancer (14) – this gap of knowledge is pronounced. HGSC is often diagnosed at advanced stages, has poor response to current immunotherapies (15,16), and is prone to chemoresistance, resulting in 5-year survival rate below 50% (14). Underscoring the need to elucidate the clinically relevant barriers that prevent anti-tumor immunity in HGSC, it is well known that, despite poor response to immunotherapies, abundant tumor infiltrating lymphocytes (TILs) are a robust prognostic marker of better clinical outcomes in HGSC patients (17,18). The genetic properties of HGSC have been thoroughly characterized (19–22) – demonstrating nearly ubiquitous *TP53* mutations, massive copy number alterations (CNA), along with mutation in homologous recombination genes such as *BRCA1* and *BRCA2* – and recent single-cell studies provided important resources and insights by characterizing the cellular properties of HGSC in different anatomical sites and genetic backgrounds (6,23). Yet, the molecular and cellular modalities that promote or suppress immune recruitment and infiltration in HGSC patients remain elusive.

Here, we applied high-plex image-based spatial transcriptomics (ST) with subcellular resolution to more than 1.36 million cells across 95 HGSC tumors. Our data demonstrates that effector T cells and NK (T/NK) cells are rarely retained in the tumor stroma. However, T/NK cell infiltration into the tumor parenchyma is skewed towards subsets of transcriptionally distinct malignant cells, leaving other malignant areas immune deserted. This, together with high-content CRISPR screens, revealed malignant cell *transcriptional* immunogenicity that is repressed by copy number alterations and can be de-repressed by an array of functionally diverse gene knockouts to sensitize ovarian cancer cells to T and NK cell cytotoxicity. Taken together, our study provides a molecular map of HGSC tumor tissue spatial organization in patients, delineates generalizable principles that predict lymphocyte location and state, and, through integration of spatial and perturbational maps, identified novel targets that reprogram tubo-ovarian malignant cell states as an avenue to unleash anti-tumor immune responses in this aggressive disease.

## RESULTS

### Single cell spatial transcriptomics mapping of tubo-ovarian high-grade serous carcinoma

To spatially map HGSC in the setting of metastatic disease, we applied *in situ* imaging with high-plex RNA detection at the single cell resolution to 95 HGSC tumors from a total of 60 patients and 136 tissue sections, yielding a total of 1,365,244 high quality single cells’ spatial transcriptomics profiles (**Figure 1a, Table S1**). Tumor sections were obtained from the adnexa (ovaries/fallopian tube, n = 73), and/or omentum (n = 63), with 37 patient-matched pairs of adnexal and omental tumors. All tumor tissue sections were obtained from debulking surgeries in either the treatment naïve (n = 66) or neoadjuvant chemotherapy treatment (n = 70) setting, with associated patient clinical data including treatment and survival outcomes (**Figure 1a**, **Table S2**, **Methods**). For 40 patients we also obtained DNA sequencing data spanning a 648-gene panel (**Figure 1b**, **Table S1**, **Methods**), focused on actionable single nucleotide variations (SNV), somatic copy number alterations (CNA), chromosomal rearrangements, and tumor mutational burden (TMB), providing a basis to probe the connection between tissue structure and somatic genetic aberrations.

**Figure 1.**
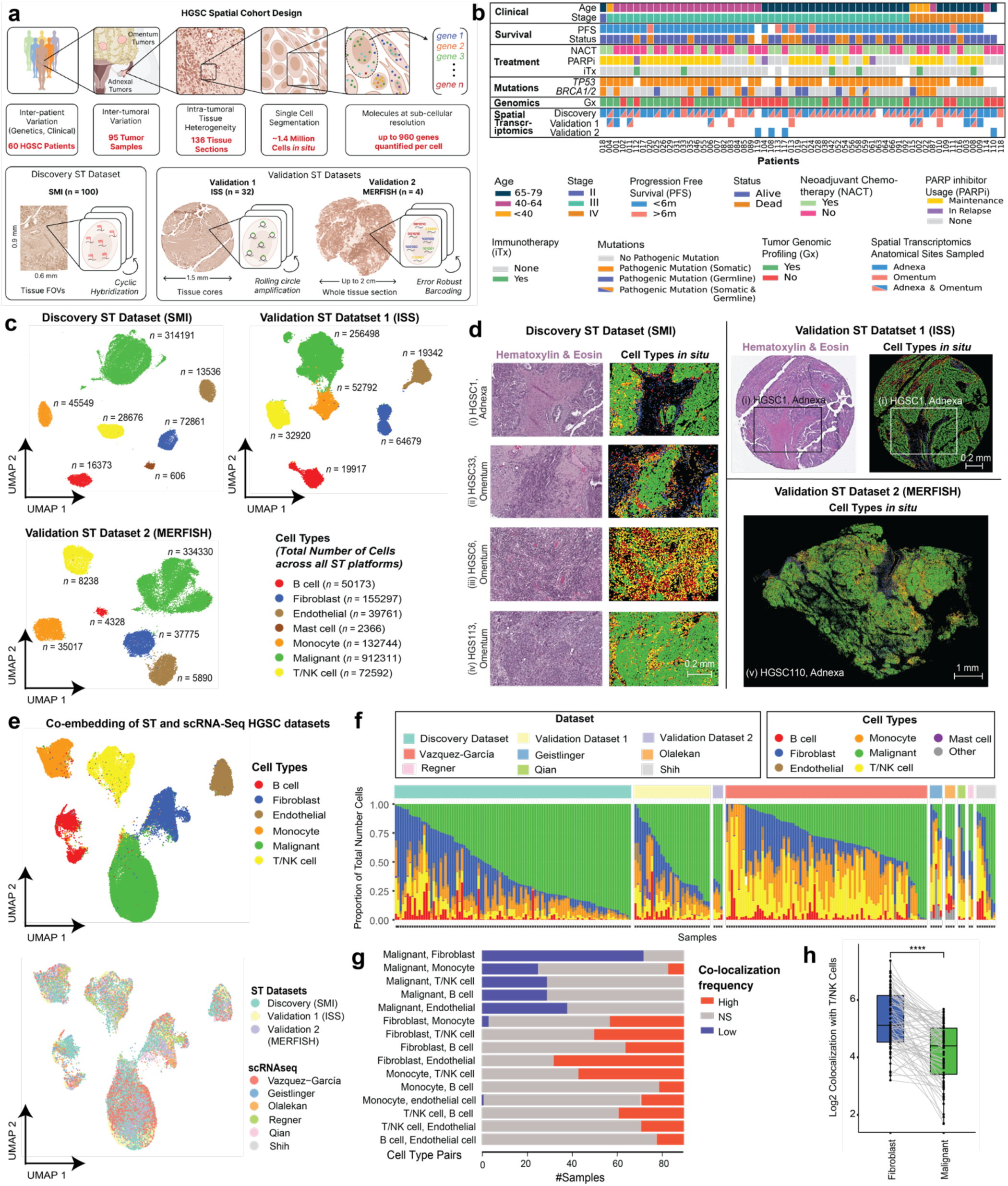
Single cell spatial transcriptomics (ST) mapping of HGSC. **(a)** Overview of the ST cohort, collected across three platforms: SMI (discovery dataset), ISS (validation dataset 1), and MERFISH (validation dataset 2); *n* denotes the number of tissue sections profiled. (**b)** Clinical annotations of the patients and samples included in the cohort. (**c)** Uniform Manifold Approximation and Projection (UMAP) embedding of cell transcriptomes from the discovery dataset (top left), validation dataset 1 (top right), and validation dataset 2 (bottom left). Cells are colored according to their cell type annotations. *n* denotes number of cells with each cell type annotation. (**d)** Representative ST images (right) and corresponding H&E (left, where available) depicting cell segmentations with each cell colored based its cell type annotations. (**e)** Co-embedding spatial cell transcriptomes from this study with publicly available scRNA-seq datasets (27,29–32,72,73). Unified UMAP of co-embedded cell transcriptomes is shown with cells colored by cell types (top) and dataset (bottom). (**f)** Cell type composition (y axis) per sample (x axis) from this study and in publicly available scRNA-seq HGSC cohorts(27,29–32,72,73). (**g)** Pairwise co-localization analysis: the number of samples (x axis) where each pair of cell types (y axis) <u>^1–3^</u>shows significantly (BH FDR < 0.05, hypergeometric test) higher (red), lower (blue), or expected (grey) colocalization frequencies compared to those expected by random. (**h)** Log_2_ Co-localization Quotient (CLQ, y-axis) of T/NK cells with fibroblasts (blue, x axis) and T/NK cells with malignant cells (green, x axis) in each tissue section from the discovery dataset (*\*\*\*\*p* < 1*10^-4^, paired Wilcoxon rank sum test). Light grey lines connect paired fibroblasts and malignant cells within each tissue section. Boxplots middle line: median; box edges: 25^th^ and 75^th^ percentiles; whiskers: most extreme points that do not exceed ± IQR x 1.5; further outliers are marked individually.

The spatial data was collected using three spatial transcriptomic (ST) platforms, allowing rigorous cross-platform validation of these recently developed technologies. A discovery ST dataset spanning 100 tissue sections was profiled with Spatial Molecular Imaging (SMI) (24), allowing *in situ* image-based quantification of 960 genes with subcellular resolution (n = 100, formalin-fixed paraffin-embedded (FFPE) tissue sections). For comparison and validation, *in situ* sequencing (ISS via Xenium(25); n = 32, FFPE tissue sections) and MERFISH (26) (Multiplexed Error-Robust Fluorescence In Situ Hybridization, n = 4, fresh-frozen tissue sections) were applied to profile 280 genes and 140 genes, respectively (**Methods, Table S1b**).

Applying a recursive clustering-based cell type annotation procedure (**Methods**) on processed gene expression profiles (**Figure S1a-d, Methods, Supplementary Information**) we identified malignant cells (n = 314,191), T and NK cells (T/NK, n = 28,676), B cells (n = 16,373), monocytes (n = 45,549), mast cells (n = 606), fibroblasts/stromal cells (n = 72,861), and endothelial cells (n = 13,536, **Figure 1c, Extended Data 1c,d**) in the discovery dataset (SMI). The same procedure resulted in similar annotations of the validation datasets (**Figure 1c**). T/NK cells were then further stratified to NK (n = 4,293), CD4 T (n = 6,040), CD8 T (n = 8,439), and regulatory T cells (Tregs, n = 1,905) in the discovery dataset (**Figure S1e-i, Methods, Supplementary Information**), with similar T/NK stratification results obtained in validation dataset 1 (**Figure S1j-k, Methods, Supplementary Information**).

Cell type annotations were validated in several ways. First, *de novo* cell type signatures identified based on the assigned cells recapitulate well known cell type markers (**Methods**, **Table S3**, **Figure S1c**). Second, cell type annotations are aligned with matching H&E and immunohistochemical markers (**Figure 1d**, **Figure S2a-d**). Third, for biological and technical replicate-matched tissue samples between the discovery and validation dataset 1, the cell type assignments aligned both spatially (**Figure S2e**) and compositionally (**Figure S2f**). Fourth, by integrating the HGSC spatial data with 6 publicly available single-cell RNA-sequencing (scRNA-seq) datasets (23,27–32), we generated a unified HGSC single-cell transcriptomic atlas (**Figure S2g**) and showed that cell type assignments aligned based on the unified co-embedding (**Figure 1e**) and corroborated the cell type annotations obtained independently based on the ST data alone (**Methods**). Lastly, using patient-matched CNA data, we show that only in malignant cells, the expression of 42% of the genes matches their CNAs (Benjamini-Hochberg False Discovery Rate (BH FDR) < 0.05, mixed-effects test; **Methods, Figure S3a**).

As expected, patients with higher T/NK cell abundance had improved clinical outcomes (*p* = 5.0*10^-2^, Univariate Cox regression Wald Test, *p* = 3.34*10^-3^ log-rank test, **Figure S2h**), and consistent enrichment of T/NK cells (*p* = 9.1*10^-3^, fisher test) and B cells (*p* = 8.29*10^-3^, fisher test) is observed in the omental vs. adnexal tumors. Malignant cells and fibroblasts are found to form spatially distinct compartments (i.e., the tumor parenchyma versus the stroma; **Figure 1g**), with significantly low intermixing between the two cell types (BH FDR < 0.05, hypergeometric test, **Figure 1g**), such that T/NK cells are preferentially localized in the stroma rather than within the tumor parenchyma (*p* < 1*10^-4^, paired Wilcoxon sum rank test, co-localization quotient, **Figure 1h**, **Figure S2i-k, Methods**).

Taken together, these findings demonstrate the quality of our data and validity of our processing pipelines and provide an initial mapping of the tumor organization into spatially segregated compartments (tumor parenchyma and stroma). This dataset sets the stage to probe into cellular transcriptional states to delineate multi-scale mechanisms underlying immune infiltration and evasion.

### Effector T cells preferentially infiltrate into the tumor

Using the ST cohort, we mapped the immune cell intrinsic and extrinsic factors that mark immune infiltration and exclusion. Starting with immune cell intrinsic properties, we mapped immune cell states as a function of their tumor infiltration status, defined based on proximity to malignant cells (**Methods**). Unsupervised embedding and clustering using single cell expression profiles alone, without any spatial information provided, shows that immune cells residing in the malignant compartment (tumor parenchyma) are transcriptionally distinct from those that reside outside (i.e., in the fibroblast compartment (stroma), **Figure 2a)**. For each of the five immune cell subtypes robustly represented in the data (CD8 T cells, CD4 T cells, Tregs, NK cells, and monocytes), we identified tumor infiltration associated genes that are significantly (BH FDR < 0.05, mixed-effect, **Methods**) over or under-expressed as a function of proximity to malignant cells (**Figure 2b,c**, **Figure S3b**, **Table S4**).

**Figure 2.**
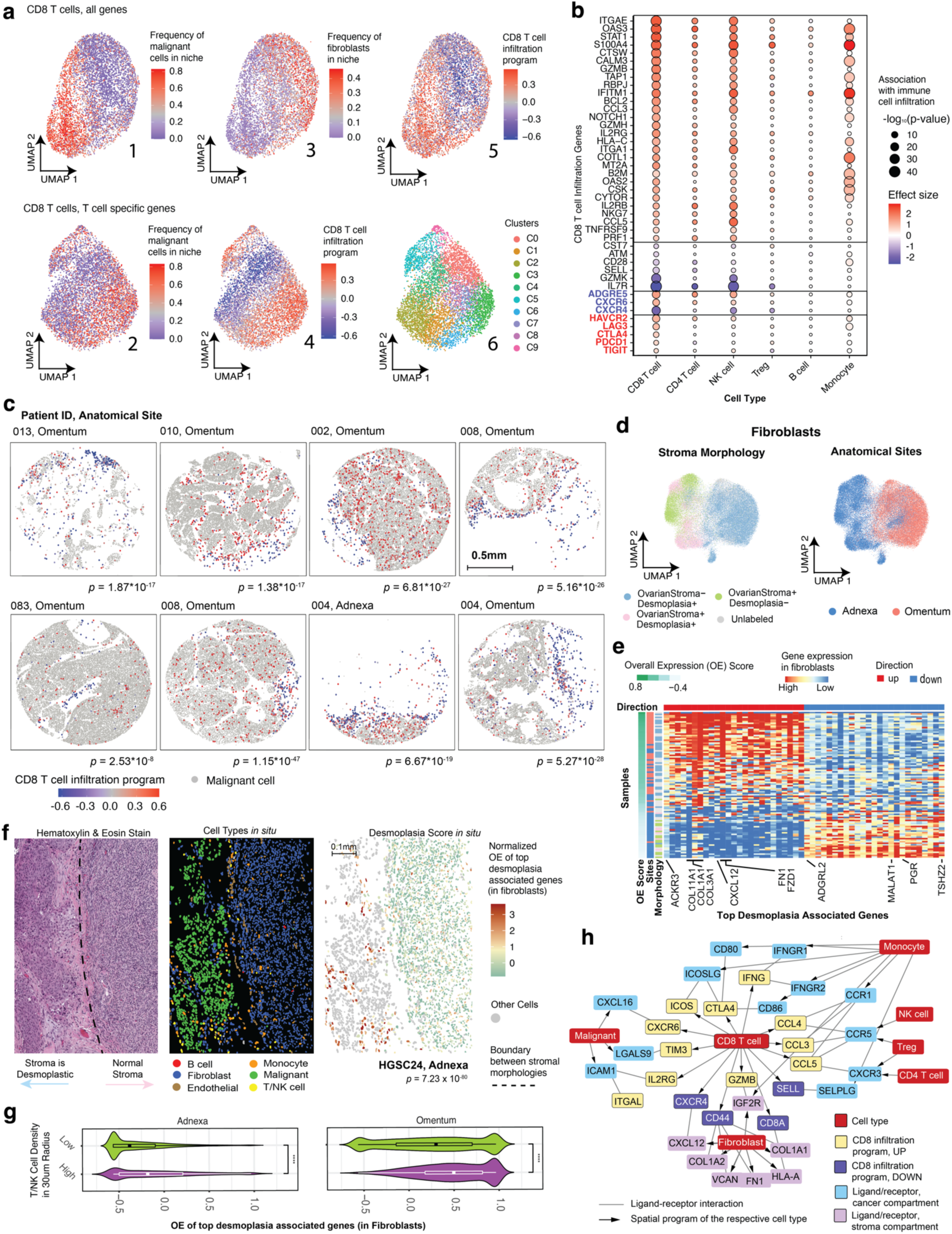
Differential immune infiltration. **(a)** UMAPs of CD8 T cells (discovery cohort) derived when considering all genes (top) or only T cell specific genes (bottom, for further confirmation). Cells are colored according to the frequency of malignant cells (1, 2) or fibroblasts (3) in the T cell microenvironment (**Methods**), the overall expression (OE) of the CD8 T cell infiltration program (4, 5) or their *k*-Nearest Neighbor cluster (6). (**b)** CD8 T cell tumor infiltration program, showing the association (*p*-value and effect size) of each gene (row) with infiltration status, when considering only specific immune cell subsets (columns). (**c)** Representative ST images from validation dataset 1 depicting the CD8 T cell tumor infiltration program identified in the discovery dataset. Malignant cells are in grey, CD8 T cells are colored according to the Overall Expression (OE) of the infiltration program identified in the discovery dataset (color bar). The respective *p*-values denote per tissue section if the OE of the CD8 T cell infiltration program is significantly higher in CD8 T cells with a high (above median) vs. low (below median) abundance of malignant cells within a radius of 30μm (one-sided *t*-test). (**d)** UMAP embedding of fibroblast cell ST profiles colored by stromal morphology (left) and anatomical site (right) annotations. **(e)** Average gene expression (z-score, red/blue top middle color bar) of the top 50 desmoplasia associated genes (columns) across fibroblasts in each sample (rows), sorted by overall expression score of the 50 genes (**Methods**, left color bar), and labeled by their anatomical site (middle color bar) and stromal morphology annotation (right color bar). (**f**) Representative tissue section (HGSC24, adnexa, discovery dataset) wherein the desmoplasia associated genes capture intra-tumoral differential stromal morphology (*p*-value = 7.23*10^-80^, Wilcoxon rank sum test). Hematoxylin & Eosin Stain (left), Cell Types *in situ* (middle), and cells plotted *in situ* with fibroblasts colored according to the overall expression (OE) of desmoplasia associated genes and all other cells in grey. (**g**) OE score of top 50 desmoplasia associated genes per fibroblast (x axis) as a function of T/NK cell density within a 30-μm radius (y axis) in the adnexa (left) and in the omentum (right). (**h)** Ligand-receptor interactions (lines) consisting of genes from the CD8 T cell infiltration program (up-regulated in yellow, down-regulated in dark purple) and their respective ligand/receptor in the cancer compartment (light blue, i.e., tumor infiltrating programs of other immune cells or genes specific to cancer cells in T cell rich areas) and stroma compartment (light purple, i.e., genes specific to fibroblasts in T cell rich areas). The arrows connect each gene to the cell type where it was found to mark the respective spatial pattern, namely, tumor infiltration in immune cells, and co-localization with T/NK cells in the non-immune (fibroblast or malignant) cells.

The CD8 T cell infiltration program demonstrates that effector and exhausted CD8 T cells are rarely excluded and frequently co-localize with malignant cells (*p* = 3.24*10^-53^, mixed effects). Tumor infiltrating CD8 T cells are characterized (BH FDR < 0.05, mixed-effects) by effector cytotoxicity genes (e.g., *GZMB* and *PRF1*) and exhaustion markers (*CTLA4*, *PD1, TIM3*), as well as the pan-cancer exhaustion marker *CXCR6* (33–35) (**Figure 2b,c**). CD8 T cells that are distant from malignant cells are characterized (BH FDR < 0.05, mixed effects) by naïve and memory T cell markers (*IL7R*, *SELL*), overexpress the chemokine receptor *CXCR4* (**Figure 2b**), and reside in the stroma, next to fibroblasts (**Figure 2a**). Expanding the CD8 T cell infiltration program to whole-transcriptome level based on scRNA-seq data (23) (**Methods**; **Table S4**) identified *TCF7* – a central regulator of naïve and resting T cells (34,35) that directly represses *CXCR6* expression – as one of the top gene negatively associated with a CD8 T cell infiltration (*p* < 1*10^-16^, *rs* = 0.23, Spearman correlation).

To investigate the role of the stroma in retaining naïve and memory T cells whilst permitting effector T cells to infiltrate the tumor parenchyma, we integrated sample-matched H&E stains independently annotated by a gynecologic pathologist (**Figure S3c**), with ST data. Analyzing these data with unsupervised embedding (**Figure 2d**) and non-linear classifiers (**Methods**) revealed two fibroblast subsets, one marking normal adnexal stroma and the other marking desmoplasia (**Figure 2d-f**, **Extended Data Fig 3d-f** i.e., a neoplasia-associated alteration in fibroblasts and extracellular matrix with distinct tissue morphology (36–40)), which we find to be more prevalent in the omentum (**Figure 2e, Figure S3e-j**). As expected (41,42), desmoplastic fibroblasts overexpress collagen fibril organization and extracellular matrix genes (*p* < 1*10^-2^, permutation test, **Figure S3d, Table S5)**, but also upregulate *CXCL12* (the cognate ligand of *CXCR4*, overexpressed in naïve/memory T cells, **Figure 2b**) and are associated with T/NK rich niches (**Figure 2g**. *p* < 1*10^-4^, mixed effects).

To systematically map spatially dependent multicellular circuits we identify all the ligand-receptor pairs that show significant (BH FDR < 0.05, mixed effects) spatial co-expression across cell types (**Methods**), revealing suppressive ligand-receptor interactions in the malignant compartment (e.g., CD80/CD86:CTLA4, CD8 T cell:monocyte; TIM3:LAGLS9, CD8 T cell:malignant cell) and CD8 T cell mediated chemoattraction of other immune cells via *CCL2* and *CCL5*. Co-localization of *CXCR6:CXCL16* (CD8 T cell:malignant cell) and *CXCR4:CXCL12* (CD8 T cell: fibroblasts) mark chemoattraction cell-cell interactions of infiltrating and excluded CD8 T cells, respectively (**Figure 2h**, BH FDR < 1*10^-10^, mixed effects test).

Collectively, these findings demonstrate a differential infiltration process wherein naïve/memory T cells primarily co-localize with the stroma, whereas effector/exhausted T cells reside primarily in the malignant compartment of the tumor (**Figure 2a-c**). While the data suggests that the stroma is not playing a major role in suppressing or trapping effector T cells (43) in HGSC patients, we find that Tumor Infiltrating Lymphocytes (TILs) are spatially segregated within the tumor parenchyma itself, as described next.

### Tumor infiltrating lymphocytes preferentially co-localize with a transcriptionally defined subset of malignant cells

Mapping the spatial distributions of Tumor Infiltrating Lymphocytes (TILs, defined here as both T cells and NK cells) revealed that TILs preferentially co-localize with a subset of malignant cells (**Methods**, **Figure 3a-c**, **Table S6**). Although malignant cell states are highly patient-specific (**Figure S4a)** and vary also within patients (**Figure S4b-d**), the connection between TIL location and malignant cell gene expression appeared repeatedly across the heterogenous tumors in our cohort (**Figure 3**, **Figure S4e-i**).

**Figure 3.**
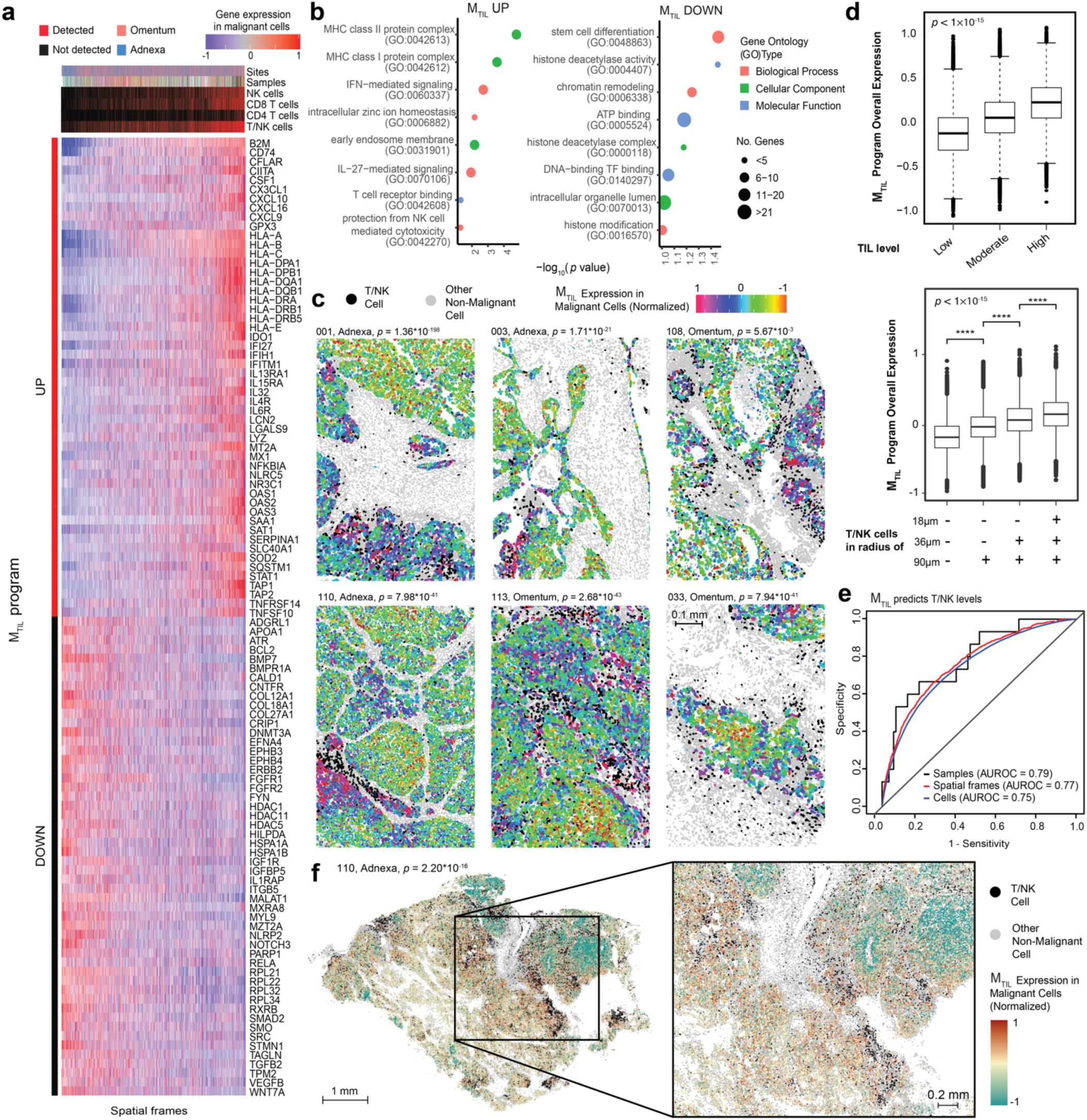
T/NK cells preferentially co-localize with a transcriptionally distinct subset of malignant cells. **(a)** Heatmap of M_TIL_ genes. Average expression (z score, red/blue color bar) of the M_TIL_ genes (rows) across spatial frames (columns), sorted by M_TIL_ overall expression (OE), and labeled by (color bar from top to bottom): anatomical site, sample ID, the detection of different T/NK cell subsets. (**b)** M_TIL_ gene ontology enrichment analysis. (**c)** Spatial distribution of T/NK cells (black) and M_TIL_ OE in malignant cells (color bar, top right) shown in representative tumor tissue sections from six different patients and anatomical sites; other non-malignant cell types are colored grey. *p*-values denote if M_TIL_ OE is significantly (one-sided t-test) higher in frames with high vs. low T/NK abundance (defined based on the median level) in the respective tissue section. (**d)** M_TIL_ OE (y axis) in malignant cells, stratified based on the relative abundance of T/NK cells in their surroundings (top) and the presence of T/NK cells at different distances (bottom). Middle line: median; box edges: 25^th^ and 75^th^ percentiles; whiskers: most extreme points that do not exceed ± IQR x 1.5; further outliers are marked individually. \*\*\*\**p* < 1*10^-4^, mixed effects (**Methods**). (**e)** ROC curves obtained for cross-validated Support Vector Machine classifier using M_TIL_ expression in malignant cells to predict T/NK cell levels, at the sample (black), spatial frames (red) and single cell levels (blue). Abbreviations: AUROC = area under the ROC curve. (**f)** Single cells visualized *in situ* in one representative whole tissue section from validation dataset 2 (left), juxtaposed with magnified region (right). T/NK cells are in black, malignant cells are colored via normalized M_TIL_ overall expression in the color bar, and non-malignant cells are in grey (M_TIL_ expression in TIL-high versus TIL-low niches, *p* = 2.2*10^-16^, Wilcoxon rank sum test).

Formulating these findings, we identified a **<u>M</u>**alignant **<u>T</u>**ranscriptional program that robustly marks the presence of **<u>I</u>**nfiltrating **<u>L</u>**ymphocytes, denoted as MTIL (**Figure 3a**, **Table S6**). The program consists of 100 up- and 100 down-regulated genes whose expression in malignant cells is significantly (BH FDR < 0.05, mixed-effects test) positively (MTIL-up) and negatively (MTIL-down) correlated with and predictive of T/NK cell infiltration (**Figure 3d,e**). MTIL overall expression in malignant cells (**Methods**) reflects both inter-and intra-sample variation in TIL levels (**Figure 3d,e**), irrespective of anatomical site (*p* < 1*10^-30^, mixed effects test; **Figure S4g**). MTIL continuously increases as a function of T/NK cell abundance and proximity (**Figure 3d**), also when stratifying the T/NK population into its respective cell subtypes (**Figure S4h**) and in validation datasets (**Figure 3f, Figure S4i**, *p* = 2.20*10^-16^). Likewise, an independent scRNA-seq dataset(44) demonstrates that the MTIL program expression in malignant cells is highest in tumors annotated as “infiltrated”, moderate in tumors annotated as “excluded”, and lowest in tumors annotated as “immune desert” (**Figure S4j**).

Gene set enrichment analyses demonstrate the connection between MTIL and immune evasion(45–50). MTIL-up includes chemokines (e.g., *CCL5, CXCL10, CXCL9,* and *CXCL16* the cognate ligand to *CXCR6*), and oxidative stress genes (e.g., *GPX3, SOD2*, **Figure 3a,b**), and is enriched with multiple immune response genes, including antigen presentation (e.g., *B2M, CIITA, HLA-A/B/C*), interferon gamma response genes (e.g., *IDO1, IFI27, IFIH1, OAS1/2/3, JAK1, STAT1*), and cell adhesion molecules (e.g., *ICAM1, ITGAV, ITGB2*; BH FDR = 1.91*10^-9^, 2.86*10^-10^, 4.59*10^-2^, respectively, hypergeometric test, **Figure 3b**, **Table S6b**). MTIL-up also includes immune suppression genes, most notable is *LGALS9*, encoding for galectin 9 – the ligand of the immune checkpoint TIM3 (i.e., *HAVCR2*), which is upregulated in the infiltrating T/NK cells (**Figure 2h**). MTIL-down reflects diverse processes including Wnt signaling (e.g., *CTNNB1, FZD3/4/6, SMO, FGFR2, WNT7A*), epigenetic regulation (*DNMT3A*, *HDAC1/11/4/5)*, as well as genes involved in insulin signaling (e.g., *IGFR1*, *IGFBP5*) and cell differentiation (BH FDR < 0.05, hypergeometric test; **Figure 3b**, **Table S6**).

The majority of the MTIL genes have never been described in the context of HGSC immune evasion. However, supporting its role in this context, a collection of CRISPR screens(45–49) assembled and analyzed here shows that MTIL-up is enriched with genes that sensitize cancer cells to immune mediated selection pressures (including *ICAM1, JAK1, NLRC5, SOD2, STAT1*, *p* = 1.82*10^-4^, hypergeometric test), while MTIL-down includes genes with desensitizing effects (*BCL2, FGFR1, HDAC1, HDAC5, ITGB5,* and *RELA*).

Given these findings, we turned to examine if MTIL repression is linked to somatic genomic aberrations as a genetic basis driving immune exclusion and tolerance.

### Copy number aberrations as repressors of the MTIL program and drivers of immune exclusion

Our cohort and independent genomic data suggest that somatic genetic variation intrinsically regulates the MTIL program and in turn impacts T/NK cell influx and non-uniform spread (**Figure 4a-c**).

**Figure 4.**
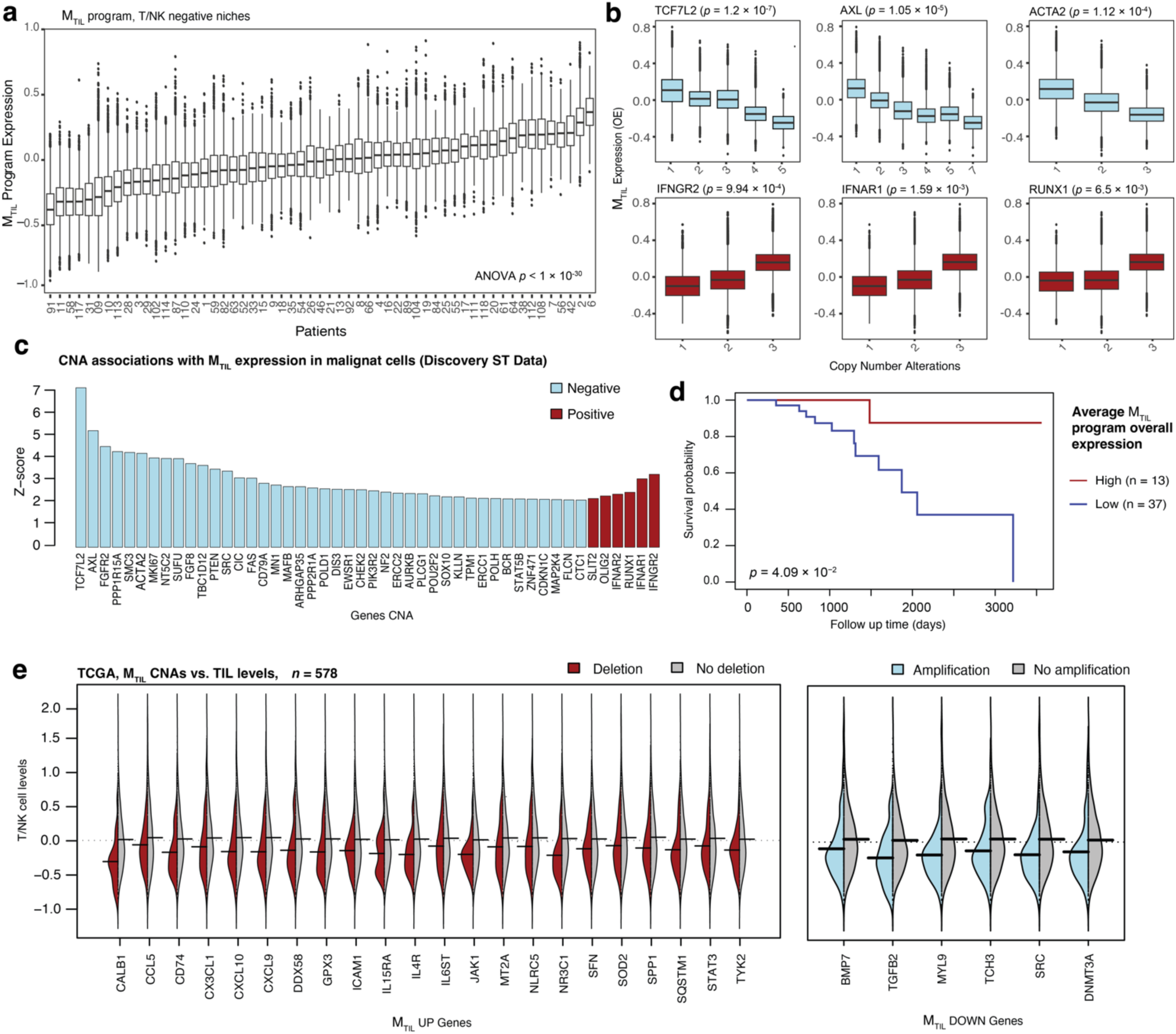
Copy number alterations associated with M_TIL_ and T/NK levels. **(a)** M_TIL_ Overall Expression (OE) in malignant cells (y axis) residing in spatial frames where T/NK cells were not detected, stratified by patients (x axis). (**b)** M_TIL_ OE in malignant cells (y axis), stratified by somatic copy number of the respective gene (x axis) based on patient-matched bulk tumor genomic profiles. (**c)** Top CNAs showing a significant (BH FDR < 0.05, mixed-effects; **Methods**) positive (red) or negative (light blue) association with M_TIL_ OE in malignant cells in the discovery spatial cohort. (**d)** Kaplan Meier Survival curves depicting differential survival probability (y axis) as a function of average M_TIL_ OE in the malignant cells of each patient (log rank test *p* = 4.09*10^-2^) (**e)** Deletion (red) of M_TIL_-up genes and amplification (light blue) of M_TIL_-down genes (x axis) are signifcantly (BH FDR < 0.05, one side t-test) associated with low T/NK levels (y axis; inferred based on gene expression of T/NK cell signatures) in an independent TCGA HGSC cohort of 578 patients (20). Grey distribution depicts the T/NK levels in tumors without the respective genomic abberation.

First, MTIL overall expression varies across HGSC patients and is associated with improved overall survival (*p* = 7.2*10^-2^, Univariate Cox regression Wald Test, *p* = 4.1*10^-2^ log-rank test, based on mean expression in the malignant cells, **Figure 4d**, **Methods**). Specifically, MTIL inter-patient variation supersedes its intra-tumoral variation, as observed also after regressing out the impact of the TME compositions, or when considering only malignant cells in TIL deprived environments (**Figure 4a**, *p* < 1*10^-30^, ANOVA test).

Second, aligned with the finding that malignant cell transcriptomes are tightly associated with CNAs in *cis* (**Figure S3a**), MTIL expression strongly correlates with the copy number of multiple genes in our cohort, the top ones being *IFNGR2* and *IFNAR1* (positively correlated) and *TCF7L2*, *FGFR2*, and *AXL* (negatively correlated, *p* < 5*10^-3^, mixed effects, **Methods**, **Figure 4b-c**).

Third, CNAs of MTIL genes are predictive of TIL abundance scores (**Methods**) in an independent TCGA cohort of 578 HGSC tumors(20) (AUROC = 0.82, on unseen test samples, supervised vector machines (SVM) model, **Methods**), where tumors with amplification of MTIL-down genes (e.g., *DNMT3A, FZD3, MYL9, SRC*, and *TGFB2*) or deletion of MTIL-up genes (e.g., *CX3CL1*, *CXCL10*, *CXCL9, ICAM1, GPX3*, *NR3C1*) have significantly lower TIL abundance scores compared to tumors without these copy number changes (BH FDR < 5*10^-3^, one-sided t-test, **Figure 4e**).

These findings propose a genetic basis to immune evasion and tolerance in HGSC, where the transcriptional CNA-driven malignant cell states can impact cancer-TIL interactions and shape TIL recruitment. To examine this model, we turned to identify regulators controlling the MTIL program and examine their functional impact on cancer cell response and susceptibility to T/NK cell-mediated cytotoxicity.

### Genetic perturbations de-repress the MTIL program and sensitize cancer cells to T cell and NK cell cytotoxicity

To functionally probe the MTIL program genes and examine their effect on cancer cell response to lymphocyte cytotoxicity, we performed high content CRISPR knockout (KO) screens in ovarian cancer cells in monoculture and co-culture with cytotoxic lymphocytes, including T Cell Receptor (TCR)-engineered CD8^+^ T cells and NK cells. Using this approach, we sought to functionally identify and distinguish between co-regulated immune response and immune suppressive genes captured by the MTIL program (e.g., *ICAM1* and *LAGSL9*) and identify perturbations that trigger the former.

Instead of targeting only genes in the MTIL program itself, we devised a meta-analysis pipeline to identify program regulators based on available Perturb-seq datasets (**Methods**). Using four previously published Perturb-seq datasets(51–53), we identified 43 and 104 perturbations that result in significantly higher or lower expression of the program, respectively (**Figure 5**, **Table S6**, **Methods**). Demonstrating the value of this approach, it revealed a wider and more diverse set of regulators, most of which are not included in the MTIL program itself or not included in the spatial data gene panels (**Table S1b**). Negative MTIL regulators are enriched for chromatin organization (e.g., *DNMT1*, *INO80*, *TAF10*, *WDR5*), Wnt pathway, Myc targets, and immune resistance genes (45–49,54) (BH FDR < 1*10^-3^, hypergeometric test). The top negative regulator identified here is *PTPN1*, which is supported by both gene activation and inhibition (**Figure 5a,c**) experiments.

**Figure 5.**
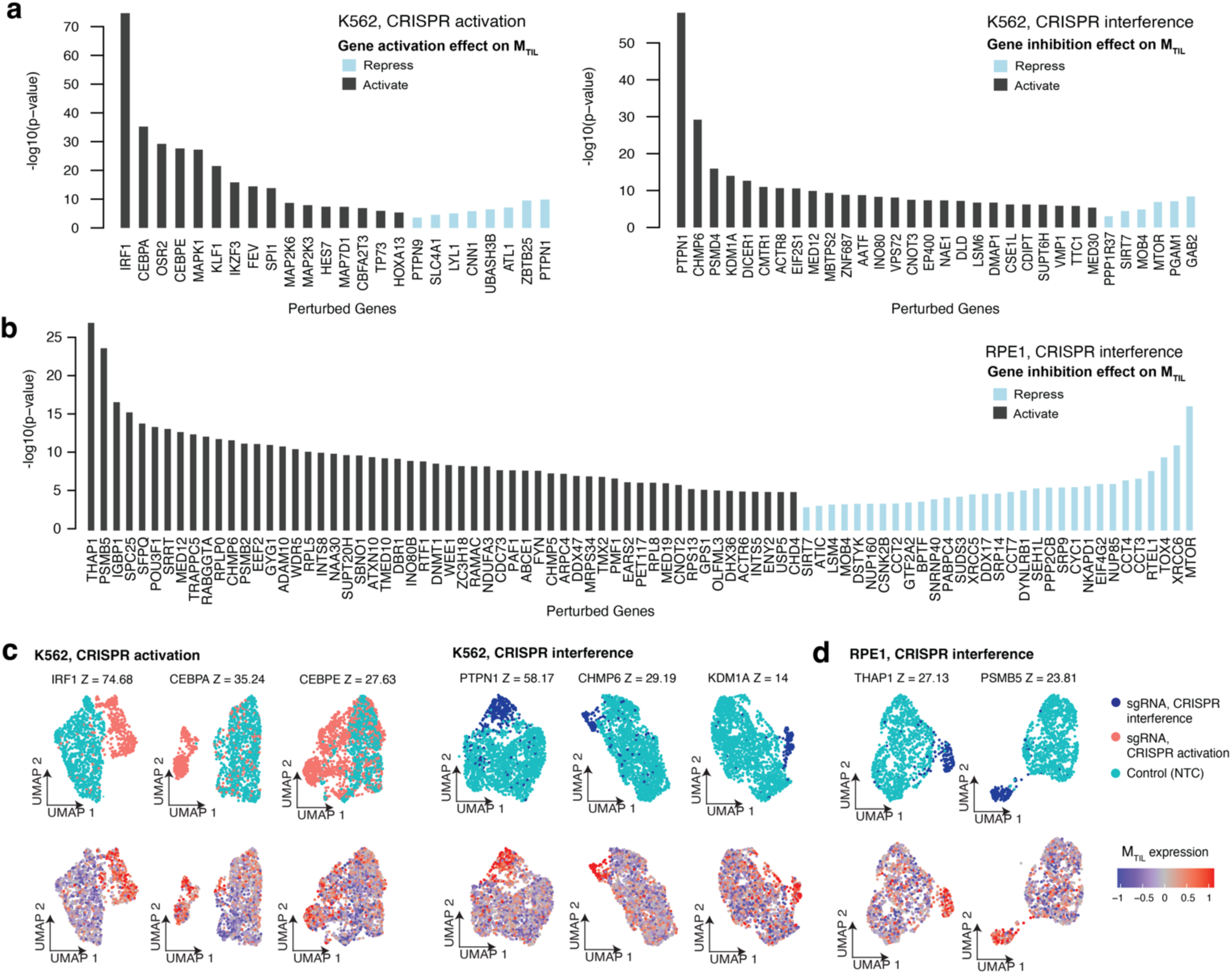
Meta-analyses of Perturb-seq datasets identifies regulators of the M_TIL_ program. **(a-b)** Differential M_TIL_ Overall Expression (OE) when comparing cells with different genetic perturbations (x axis) to cells with control sgRNAs, showing the statistical significance (two-sided *t*-test, y axis) for the top perturbations identified to repress (light blue) or activate (grey) M_TIL_ expression in (**a)** K562 (myelogenous leukemia) and (**b)** RPE1 (human retinal pigment epithelial) cell lines Perturb-seq data(51,52). (**c-d)** Representative UMAP embeddings of M_TIL_ altering perturbation: cells are labeled based on the sgRNA detected (top) and based on M_TIL_ OE (bottom) in K562 (c) and RPE1 (d) cell lines. *Z* denotes −log_10_ (*p*-value), two-sided *t*-test, comparing M_TIL_ OE in the perturbed vs. control cells.

This approach guided our design of the pooled knockout of 74 MTIL genes and regulators (**Table S7**) in ovarian cancer cells (TYK-nu cell line, **Figure 6a, Figure S5–6**). Mapping fitness upon genetic perturbations under both innate and adaptive immune selection pressures (**Figure 6a,b**, BH FDR < 0.05, MAGeCK, **Methods**) along with Perturb-seq scRNA-seq readouts in monoculture and co-culture with NK cells (**Figure 6a,c**), allowed us to identify perturbations that activate or repress the program and track subsequent effects of these perturbations on immune escape. In total we profiled 18,585 high quality single cell transcriptomes, each assigned to an ovarian cancer cell with a single sgRNA confidently identified, and a median of 4,251 genes detected per cell (**Figure 6c**, **Figure S7a**). Differentially expressed genes were identified for each gene knockout across the three conditions (fisher method; **Methods**), resulting in 74 gene “perturbation signatures” (**Methods**) that were then used to identify gene knockouts that significantly repress or activated the MTIL program, denoted as -activators and -repressors, respectively (**Figure 6d**, **Methods**).

**Figure 6.**
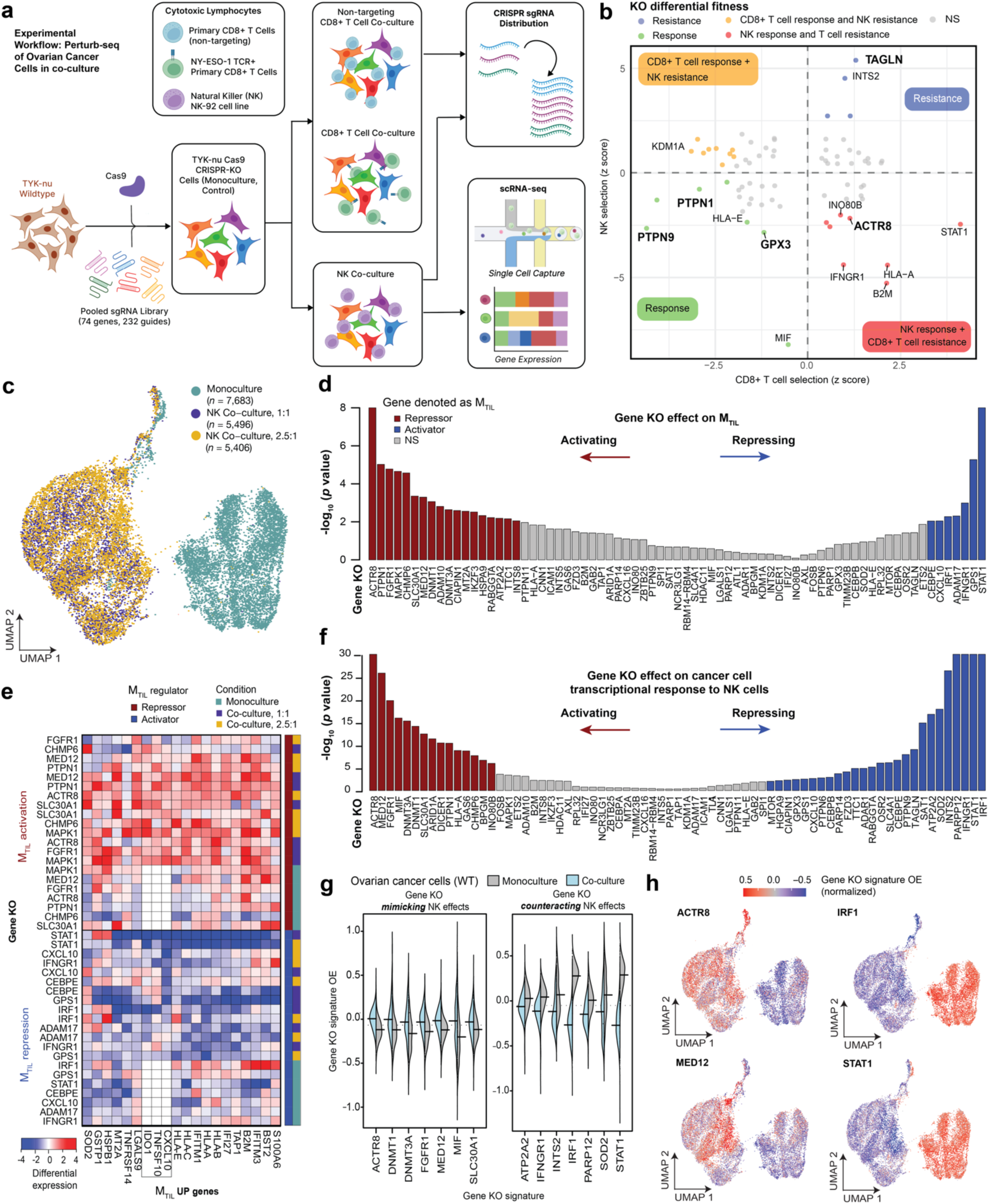
High content CRISPR screen uncovers genetic perturbations that de-repress M_TIL_ expression and sensitize ovarian cancer cells to NK and CD8 T cell cytotoxicity. **(a)** Overview of experimental design. Abbreviations: TCR = T cell receptor, KO = knockout. (**b)** Ovarian cancer cell (TYK-nu) differential fitness (MAGeCK (74)) under CD8^+^ T cell selection pressure (x axis) and NK cell selection pressure (y axis), showcasing gene KOs that confer response (green), resistance (blue) to both CD8^+^ and NK cell-mediated killing, or differential response and resistance to CD8^+^ T and NK cell-mediated killing (orange and yellow). (**c)** UMAP of single cell RNA-seq (scRNA-seq) profiles from Perturb-seq screen. Each dot corresponds to an ovarian cancer cell (TYK-nu) with one of the 232 guides confidently detected, cultured in monoculture (blue) or co-culture with NK cells in 1:1 (purple) or 2.5:1 (yellow) effector to target ratio. (**d)** Differential expression of M_TIL_ genes (fisher combined test; **Methods**) when comparing ovarian cancer cells with the respective gene KO to those with non-targeting control (NTC) sgRNAs. **(e)** Differential expression M_TIL_-up genes (columns) upon different gene KOs (rows) under different conditions (monoculture and co-culture with NK cells; vertical rightmost color bar), shown for genes identified as M_TIL_ repressors (red) or activators (blue). (**f-h)** Gene knockouts alter the cancer cell transcriptional response to NK cells. **(f)** Gene knockouts (x axis) activate (red) or repress (blue) the cancer cell transcriptional response to NK cells (t-test p-values, y axis). (**g)** Overall expression (y axis) of different gene KO signatures (x axis) in unperturbed ovarian cancer cells in monoculture (grey) and co-culture (light blue); statistical significance shown in (f). (**h)** UMAPs as in (a) with cells colored according to the overall expression of four different gene KO signatures.

Validating our hypothesis and approach, the top perturbations activating the program – *PTPN1* and *ACTR8* knockouts – sensitize malignant cells to T/NK cell cytotoxicity (**Figure 6b,d-e, Figure S7b**), while the top perturbations that repress the program, *IFNGR1* and *STAT1* knockouts, allow ovarian cancer cells to resist T cell mediated killing (**Figure 6b,d-e, Figure S7b**). Knockout of *ACTR8* and *PTPN1*, as well as other top repressors *FGFR1*, *MAPK1*, and *MED12* were found to sensitize cancer cells to immune elimination also in based on data from previous *in vivo* CRISPR screens(45–49). Moreover, we find that knockout of MTIL repressors (*ACTR8, DNMT1, FGFR1, PTPN1, MED12,* and *MIF*) mimics and amplifies the transcriptional responses to NK cells, while knockout of MTIL activators, as *STAT1*, *IFNGR1*, *INTS2*, *IRF1*, *PARP12* and others, represses and counteracts the transcriptional response to NK cells (**Figure 6f-h**, **Extended Data 7c-f**). Lastly, knockout of specific genes within the program, including *GPX3* and *TAGLN* show substantial impact on the ovarian cancer cell susceptibility to NK mediated killing (**Figure 6b**), demonstrating both global and gene-specific effects.

Taken together, coupling HGSC spatial tumor organization with multimodal functional probing, identified new and clinically relevant targets to sensitize ovarian cancer cells to innate and adaptive cytotoxic lymphocytes and demonstrated the role of cancer cell intrinsic transcriptional dysregulation as an important driver dictating the outcomes of the malignant-T/NK cell interplay.

## DISCUSSION

Our study maps the tumor tissue landscape in HGSC patients and reveals generalizable principles of tissue organization that dictate lymphocyte location and state within these aggressive and genetically unstable tumors. It uncovers a profound connection between somatic genetic aberrations, malignant transcriptional dysregulation, and immune evasion at the cellular and tissue level, providing a new perspective to the barriers preventing the anti-tumor immune response in HGSC patients and new leads to derepress HGSC cancer immunogenicity.

Innate and adaptive cytotoxic lymphocytes (CTLs) have a substantial effect on cancer cell transcriptome (**Figure 6c**). As shown here, genetic dysregulation that prevents this transcriptional response can have significant effects on cancer cell susceptibility to immune elimination even in the highly controlled co-cultures as those used here, where CTLs are already primed and activated, and spatial segregation is unlikely to occur. These effects can be amplified in the context of *in vivo* cancer-immune co-evolution where immune tolerance is reinforced due to positive feedback loops across cells. Indeed, immune checkpoint blockade and other immunotherapies have shown modest effects in tumors with low TIL levels at baseline (17,55). The data shown here proposes that this may not be only due to immune exclusion per se, but also due to *cancer intrinsic* differences between TIL-rich and TIL-deprive tumors that protect malignant cells even in the presence of targeting CTLs. Our findings and approach open new directions for further investigation of the genetic basis of tumor immune evasion through the lens of spatial organization and put forward a framework to design targeted strategies to counteract or bypass these resistant mechanisms.

More generally, as more spatial datasets become available, there is a growing need to use this rich information to delineate new drivers of complex multicellular processes and phenotypes. Here we show the value of mapping spatial cell states to genetic information across individuals and to design perturbational screens with single cell readouts. Importantly, we show that using existing Perturb-seq datasets to identify latent regulators of gene expression programs is critical and provides a data-driven framework to uncover regulators that are not necessarily included in the program itself. As more Perturb-seq datasets, as the one generated here, become available across a more diverse range of cell types and conditions, it will be possible to use this information more effectively to extrapolate from one context to another with increasing accuracy (56,57).

The key findings from our study can fuel new lines of investigation towards new clinical interventions in HGSC patients. We anticipate that the detailed mapping of HGSC tumors provided here will help inform the design of new T/NK cell engineering strategies to reach better control of cell delivery and location in a more precise manner that is aligned with the tumor cellular and molecular structure in patients. Our findings demonstrate that the stroma forms a differential “filter” that supports differential occupancy of effector T cells in the malignant compartment – this calls for dynamic tracking of tumor reactive T/NK cells across non-tumor sites (i.e., in the circulation and lymph nodes) and within the tumor to help elucidate this process and examine if T cells can also egress back to the stroma cells to avoid or reverse exhaustion(58) and how to best leverage, as opposed to eliminate or target, the stroma.

Our data provides new leads to target HGSC resistance, including epigenetic regulators (e.g., *ACTR8* and *MED12*), fibroblast growth factor receptors (*FGFR1/2*), *GPX3*, and *PTPN1*. *PTPN1*, which we found to be one of the most potent MTIL repressors, provides pre-clinical rationale to test new *PTPN1/N2* inhibitors (NCT04777994, NCT04417465, phase I clinical trials)(59–64) in HGSC patients, and demonstrates a connection between immune evasion, insulin resistance, and type 2 diabetes. *PTPN1* is a negative regulator of insulin and leptin signaling (65) that has been an attractive drug target for treatment of type 2 diabetes and obesity (66–69). *PTPN1*’s protein product PTP1b is inactivated by oxidation(70), which may explain MTIL activation under oxidative stress (as indicated by the up regulation of *GPX3* and *SOD2*). Further supporting the connection to insulin resistance, *TCF7L2*, which we identified as a top gene amplification associated with the repression of the MTIL program in the HGSC cohort (**Figure 5b-c**) harbors the most significant SNP associated with type 2 diabetes risk (71).

Taken together, this integrative study provides a blueprint to functionally map and probe the molecular landscape of multicellular interplay in complex biological tissues and reveals unrecognized spatial, molecular, and genetic regulation of immune escape in HGSC, opening new avenues to activate targeted immune responses in this aggressive disease.

## METHODS

### Human tumor specimen collection

For the discovery (SMI) and validation 1 (ISS) spatial cohorts, all tumor tissues were archival clinical formalin fixed paraffin embedded (FFPE) tumor tissues, retrospectively procured from archival storage under Institutional Review Board (IRB) approved protocol (#44615). In patients with both adnexal and omental tumors available for study, tumor blocks from both sites were selected by an expert gynecologic pathologist (B.E.H) using histopathologic review of the associated H&E slides. HGSC diagnosis was confirmed in all cases. Tumor content as well as tissue quality and preservation were assessed for inclusion in the study. For the validation 2 (MERFISH) ST dataset, fresh HGSC tumors were collected at the time of surgery by Stanford Tissue Procurement Shared Resource facility with the appropriate written informed consent and institutional IRB approval (#11977). Samples were flash frozen and stored at −80℃ until requested for this study. Samples were embedded in optimal cutting temperature (OCT). Sections were generated using a cryostat and stained with H&E, which were reviewed by an expert gynecologic pathologist (B.E.H.) to confirm the diagnosis, quality, and tumor content. Summary statistics of tissue sections, tumors, and patients profiled are available in **Table S1a**. Annotations at the patient level and tissue level are provided in **Figure 1b** and **Table S2**.

### Bulk tumor tissue Next Generation Sequencing (NGS)

HGSC tumor sample selection for NGS was based on the assessment of overall tumor content by a board-certified expert pathologist (B.E.H). Solid tumor tissue was digested by proteinase K. Total nucleic acid was extracted from FFPE tissue sections using Chemagic 360 sample-specific extraction kits (Perkin Elmer). Percent tumor cellularity as a ratio of tumor to normal nuclei was verified against pathologist-derived assessment, with a minimum requirement of 20% tumor content. Macro-dissection was utilized as required to enrich specimens below the 20% threshold. Specimens that met the 20% threshold of tumor to normal nuclei were selected for DNA sequencing. DNA sequencing was subsequently performed via Tempus Labs according to the xT platform protocol (75). Additional information about NGS data generation and processing is provided in **Supplementary Information**.

### Spatial Molecular Imaging (SMI)

SMI data was generated using the CosMx SMI instrument, according to the company’s protocols, as described here and in **Supplementary Information**. CosMx pre-commercial RNA 960 gene panel was used (**Table S1b**), consisting of in situ hybridization (ISH) probes. Each reporter set contains 16 readout rounds with four different fluorophores, creating a 64-bit barcode design with Hamming distance 4 (HD4) and Hamming weight 4 (HW4) to ensure low error rates. Probe fluorescence was detected at subcellular resolution via the CosMx SMI instrument and the signal was aggregated to identify the specific RNA molecule measured in each location (24).

#### SMI tissue preparation and RNA assay

Five-micron tissue sections were cut from FFPE TMA tissue blocks and adhered onto VWR Superfrost Plus Micro Slides (VWR, 48311-703). After sectioning, the tissue sections were air-dried overnight at room temperature. Tissue preparation was performed as described in the CosMx SMI Manual Slide Preparation Manual (MAN-10159-01). Briefly, the tissues underwent deparaffinization, heat-induced epitope retrieval using a pressure cooker for 15 minutes at 100℃, and enzymatic permeabilization with 3 µg/mL digestion buffer for 30 minutes at 40℃. Subsequently, a 0.0005% working concentration of fiducials were applied to the tissue, followed by post-fixation and blocking using NHS-acetate. Finally, an overnight hybridization was performed using the pre-commercial 960 plex RNA Panel of probes. The next day, the tissues were subjected to stringent washes to eliminate any unbound probes. The tissues were stained with CosMx Nuclear Stain, CosMx Hs CD298/B2M, CosMx Hs PanCK/CD45, CosMx Hs CD3 nuclear and segmentation markers. An additional round of blocking using NHS-acetate was performed before loading on to the instrument. The slide and coverslip constitute the flow cell, which was placed within a fluidic manifold on the SMI instrument for analyte readout and morphological imaging. Analysis run on the instrument was set up using the 60 seconds per FOV pre-bleaching profile and segmentation profile for human tissue. Twenty FOVs were selected per slide resulting in a total of 100 tissue profiles.

### In Situ sequencing (ISS)

ISS was performed using 10X Genomics’ Xenium platform, according to the company’s protocols as described here and in **Supplementary Information**. In brief, 10X Genomics’ Xenium ISS technology was uses with the Xenium Human Breast Panel that consists of 280 genes (**Table S1b**). Xenium hybridization padlock probes were designed to contain two complementary sequences that hybridize to the target RNA(76). Probes also contain a third sequence encoding for a gene-specific barcode such that once the paired ends of the probe bind to the target RNA and ligate a circular DNA probe is generated for Rolling Circle Amplification (RCA). This system increases specificity and minimizes off-target signals as ligation should not occur in off-target binding events.

#### Xenium tissue preparation and RNA assay

Five-micron FFPE TMAs were sectioned onto a Xenium slide. Deparaffinization and permeabilization was performed to expose mRNA. The mRNAs were targeted by 280 probes and two negative controls: one to assess non-specific binding and the other a genomic DNA (gDNA) control to ensure signal comes from mRNA. Tissue slides were incubated overnight at 50℃ with a probe concentration of 10 nM. Following stringent washes to remove un-hybridized probes, probes were ligated for two hours at 37℃. At this step an RCA primer was annealed. Circularized probes were enzymatically amplified for one hour at 4℃ followed by two hours at 37℃ to generate multiple copies of gene-specific barcodes for each RNA binding event which increases the signal-to-noise ratio. Following washing, background fluorescence was quenched chemically to mitigate auto fluorescence that is caused by lipofuscins, elastin, collagen, red blood cells and formalin fixation (76). Slides were placed into an imaging cassette and loaded on the Xenium Analyzer instrument.

#### Image pre-processing

The Xenium Analyzer captured a Z-stack of images every cycle and in every channel. Images were processed and stitched to build a spatial map of the transcripts across the tissue section. Stitching was performed on the DAPI image, taking all the stacks from different FOVs and colors to create a single image representative of one tissue section.

### Multiplexed Error-Robust Fluorescence In Situ Hybridization (MERFISH)

Data was generated on the Vizgen Inc. platform according to the company’s protocols. A custom 140 gene panel was designed with an additional set of 50 blank negative control barcodes based on the MERFISH design that incorporates combinatorial labeling with an error-robust encoding scheme to mitigate detection errors(77).

#### Tissue Sectioning and Permeabilization

Four HGSC fresh frozen tissue samples were preserved in OCT compound and stored at −80℃ prior to sectioning. Ten-micron tissue sections were cut from the fresh frozen OCT tissue blocks and adhered onto MERSCOPE slides (Vizgen, 20400001). After sectioning, the tissue sections were fixed with 4% paraformaldehyde in 1X PBS for 15 minutes, washed three times with 1X PBS, and incubated overnight at 4℃ in 70% ethanol.

#### Cell boundary and antibody stain

Following overnight permeabilization, the tissue sections were placed in the MERSCOPE Photobleacher (Vizgen, 1010003) for four hours to quench autofluorescence.

Subsequently, the tissue samples were stained using Vizgen’s Cell Boundary Kit (Vizgen, 10400009) and blocked in blocking solution (Vizgen, 20300012) supplemented with a 1:20 dilution of Rnase inhibitor (NEB, M0314L) for one hour. The tissue sections were washed with 1X PBS and stained with the Cell Boundary Primary Staining Mix (Vizgen, 20300010) at a 1:100 dilution supplemented with a 1:20 dilution of Rnase inhibitor for one hour. After a series of washing with 1X PBS, the tissue samples were stained with the oligo conjugated secondary antibodies that were supplemented with 1:20 dilution of Rnase inhibitor for one hour. After incubation, the tissue sections were fixed with 4% paraformaldehyde in 1X PBS for 15 minutes and washed in 1X PBS.

#### Encoding Probe Hybridization

Tissue sections were washed for five minutes in Sample Prep Wash Buffer (Vizgen, 20300001) and incubated in Formamide Wash Buffer (Vizgen, 20300002) for 30 minutes at 37℃. The custom MERSCOPE Gene Panel Mix (Vizgen, 20300008) was applied to the tissues and the slides were incubated at 37℃ for 36 to 48 hours. After hybridization, the tissue sections were washed with Formamide Wash Buffer for 30 minutes at 47°C and then subsequently washed with Sample Prep Wash Buffer for two minutes.

#### Gel embedding and tissue clearing

Tissue samples were embedded in a gel solution and incubated for one minute. The gel was made from a gel embedding solution that was comprised of Gel Embedding Premix (Vizgen, 20300004), 10% ammonium persulfate (Sigma, 09913-100G), and TEMED (N,N,N’,N’-tetramethylethylenediamine, Sigma, T7024-25ML). The gel solution was then removed, and an additional gel solution was added to the top of the sample sandwiched beneath a 20 mm Gel coverslip (Vizgen, 20400003). The samples were incubated for 1.5 hours at room temperature to allow the gel solution to polymerize. The gel coverslips were prepared using RNAseZap and 70% ethanol prior to being covered with Gel Slick (VWR, 12001-812). Following the incubation, the Gel Coverslip was removed, and the samples were incubated overnight at 47℃ in clearing solution comprised of Protease K (NEB, P8107S) and Clearing Premix (Vizgen, 20300003). The tissue samples were then incubated overnight at 37℃.

#### Sample Imaging

Following the series of overnight incubations, tissue sections were washed in Sample Prep Buffer for ten minutes and incubated at room temperature for 15 minutes in DAPI and Poly T Reagent (Vizgen, 20300021). Tissue samples were then washed in Formamide Wash Buffer for ten minutes, transferred into Sample Prep Wash Buffer, and loaded onto the MERSCOPE instrument (Vizgen, 10000001) for morphological imaging and analyte readout. The appropriate fluorescently labeled probe solution was applied to the tissue, imaged, and photobleached to remove the probes for the next round.

### Cell segmentation

Cell segmentation was performed using a deep-learning based segmentation image processing algorithm, Mesmer (78) (**Figure S1a,b**) from within the DeepCell platform on raw TIFF images. The inputs for whole cell segmentation for SMI images included immunofluorescent (IF) images of DAPI and CD298/B2M for nuclear and cell membrane detection, respectively. Similarly, MERFISH whole cell image segmentation was performed with DAPI and cell membrane stains (Vizgen stain boundary kit, 10400009). Nuclear segmentation was performed for ISS images wherein the input includes DAPI IF stain.

### Gene expression quantification from spatial transcriptomics data

Preprocessed RNA *in situ* data includes RNA transcripts confidently identified for each gene and their spatial coordinates. Given this data each RNA transcript was aligned to the cell segmentation outputs described above based on its spatial coordinates. Cell count matrices, *C*, were generated by counting the number of RNA transcripts detected within the segmentation boundaries of each cell *j* for each gene *i* to yield *c*_i,j_ for entry of *C* in each ST dataset. Cell counts were converted to transcripts per million:

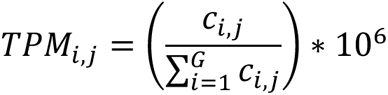

wherein *G* is the total number of genes in each ST dataset.

Expression levels were quantified as:

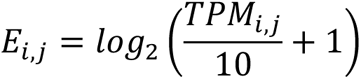

The average expression of a gene *can* across a population of N cells, denoted here as *P*, was defined as:

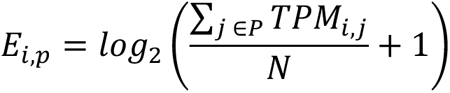

Cells with fewer than 50, 20, and 5 genes detected in the SMI, Xenium, and MERFISH data were excluded, as well as cells with exceptionally large volume (> 441 μm^2^).

Overall Expression (OE) of a gene signature was computed with additional normalizations to filter technical variation, similar to the procedure reported before (79) with some modifications as described in **Supplementary Information** “Gene set Overall Expression (OE)”.

### Cell neighborhoods and niche definition

The location of each cell was defined based on the location of its centroid. The *r*-neighborhood of a cell was defined as all the cells that reside at a distance of at most *r* μm from the cell. Spatial frames were defined by binning the tissue section FOV to 75 μm x 75 μm (i.e., 5625 μm2) sized squares, with a median number of 53 cells per frame.

### Cell type annotations

The cell type annotation procedure was applied separately for each of the three spatial datasets via an initial cell type assignment followed by an iterative subsampling procedure to obtain robust cell type assignments with confidence levels. The pipeline is described in **Supplementary Information** “Cell type annotations” and “T/NK subtype annotations” and will be provided in the study GitHub repository.

### Deriving cell type signatures

scRNA-Seq data of three HGSC cohorts was used to identify cell type specific signatures (**Table S3a**) used for cell type annotations. Preprocessed gene expression matrixes were downloaded from https://lambrechtslab.sites.vib.be/en/data-access (30,32), GSE146026 (80), and GSE173682 (29). Cell type annotations as reported in each of these studies were used. For each of the three cohorts cell type signatures (*t*-test *p*-value < 1*10^-10^ and log_2_ fold change > 0.2 for all pairwise comparisons) were derived and genes supported by at least two datasets were included in the final signatures. Cell type signatures were also derived from the discovery data set (**Table S3b**) via a similar approach with modifications described in **Supplementary Information** under “Deriving cell type signatures from spatial transcriptomics data”

### Co-embedding for a high-quality reference single cell atlas

A reference single cell atlas was generated to examine consistency across spatial and scRNA-seq cohorts and validate cell type annotations. The atlas includes three spatial datasets collected here and six scRNA-seq HGSC cohorts (23,27–32). Preprocessed gene expression matrixes were downloaded from Synapse (syn33521743 (23)), GSE118828 (27), GSE173682 (29), GSE147082 (28), GSE154600 (31), and https://lambrechtslab.sites.vib.be/en/data-access (30,32).

Tumor samples derived from other anatomical sites, other than the adnexa or omentum, were removed to match the scope of this study. For each scRNA-seq dataset, max{5000, *N_t_*} cells were subsampled from each of cell type annotations, where *N_t_* denotes the number of cells labeled with cell type annotation *t*. Each of the ST datasets were subsampled as follows. For each non-malignant cell type, max{5000, *N_t_*} high confidence cells were subsampled, where *N_t_*denotes the number of high confidence cells labeled with cell type annotation *t*. From malignant cells, max{500, *N*_*mp*_} high confidence malignant cells were sampled per patient, where {*N*_*mp*_} denotes the number of malignant cells per patient *p*.

All nine subsampled datasets were co-embedded with reciprocal principal components analysis (RPCA) using the top 30 PCs fit on each dataset, using the Seurat R Package v4 implementation (81), and then visualized with two dimensional Uniform Manifold Approximation and Projection (UMAP) (82). More details on cell type matching across publicly available datasets are available in **Supplementary Information** under “Cell Type Annotation Harmonization across Datasets”.

### Mixed effects modeling

Mixed effect models were used to capture co-dependencies and the hierarchical structure of the data, where covariates at different levels (e.g., cell, spatial frame, sample, etc.) are sampled from different distributions.

The following model was used to decompose a feature of interest *y*:

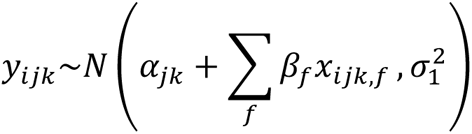

Where *y_ijk_* is the value of the feature in cell *i* in frame *j* of sample *k*, *x_ijk,f_* is a cell-type-level covariate (*e.g.*, log-transformed number of reads), and α*_jk_* is the spatial frame *j* intercept, defined as

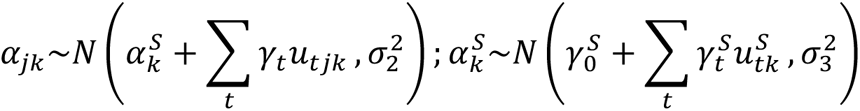

where *u_tjk_* are frame-level covariates of frame *j* in sample *k* (e.g., T cell abundance in the frame), and 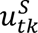 are sample-level covariates of sample *k* (e.g., sample site, treatment status). Patient-level covariates were added in the same manner as a fourth level. The lme4 (83) and lmerTest R packages (84) were used to fit the model, compute *p*-values and sum of squares (*SS*) in type II ANOVA (<u>aN</u>alysis Of <u>vA</u>riance) via the Satterthwait’’s degrees of freedom method, and identify the latent variables that maximize the posterior probability.

### Spatial transcriptional program identification

Immune infiltration programs (**Table S4**) were identified with the mixed-effect models described above using the frame-level abundance of malignant cells as a measure of the infiltration level. To prevent impact of ambient RNA, only genes that have a significantly higher expression levels (pairwise one-sided *t*-test *p*-value > 1*10^-3^) in respective cell type were considered, using pairwise *t*-tests when comparing the respective cell type to all other cell types. The CD8 T cell infiltration program was extended based on scRNA-seq data (23). Analyzing the CD8 T cells from this scRNA-Seq cohort the top 50 genes that were significantly correlated (BH FDR < 1*10^-10^, Spearman Correlation) with the OE of the CD8 T cell infiltration signature were identified (**Table S4**). Malignant TIL (MTIL) program (**Table S6**) was identified in a similar manner, using the presence of T/NK cells as a binary covariate at the frame-level. P-values were corrected for multiple hypotheses testing using the BH test, and topmost genes with FDR < 0.05 were reported.

### CNA and anatomical site analyses of spatial transcriptomics

Mixed-effect models were used to compute the association between the expression of each gene in the different cell types and the patient-matched CNA measurements obtained at the bulk tumor level. Of the 626 genes with CNA measurements, 159 were also included in the discovery dataset (SMI) panel. For each cell type and each of these 159 genes the following model was fit: tpm ∼ (1 | patient) + cna + nact + sites, where tpm denotes the expression of the gene in cells from cell type *k*, cna denote the copy number of the same gene, nact denotes treatment status, and sites denote the anatomical site. Similarly, to examine the connection between the MTIL program OE and CNAs, all 626 genes with CNA were tested with this model, considering only malignant cells from the samples with genomic profiling, with MTIL OE as the dependent variable. To derive associations of treatment status and anatomical site, a similar model was fit (tpm ∼ (1 | patients) + nact + sites) on all 960 genes in the discovery dataset.

### CNA analyses of TCGA data

The Cancer Genome Atlas (TCGA) data of Array-based Gene Expression (EXP-A) and Copy Number Somatic Mutations (CNSM) was downloaded from the International Cancer Genome Consortium (ICGC; https://dcc.icgc.org/projects). The TIL levels of each sample were computed as the overall expression of T cell signatures. Amplifications and deletions were defined as a copy number log-transformed value (“segment_mean”) above or below 0.5 and −0.5, respectively. To examine the hypothesis that repressing the MTIL-up genes drives T cell exclusion, a one-sided t-test was preformed to examine if samples with deletion in the MTIL-up genes have significantly lower TIL scores compared to all other samples. Likewise, to examine if induction of MTIL-down genes drives T cell exclusion, a one-sided t-test was preformed to examine if samples with amplifications in the MTIL-down genes have significantly lower TIL scores compared to all other samples. Support Vector Machine (SVM) classifiers were generated to predict if a tumor has a high (above median) TIL levels based on the CNA levels of all MTIL genes, using the e1071 R package.

### Perturb-seq meta-analyses and target selection

Publicly available Perturb-seq datasets were used to identify MTIL regulators and identify targets for the ovarian cancer Perturb-seq screen performed here. The Perturb-seq collections used include the following datasets. (1) large-scale CRISPR KO Perturb-seq screens in K562 and RPE1 cell lines (52). Counts data and metadata were downloaded from https://gwps.wi.mit.edu/, focusing on K562 day 8 Perturb-seq (KD8; targeting all expressed genes at day 8 after transduction) and RPE1 day 7 Perturb-seq (RD7; targeting DepMap essential genes at day 7 after transduction). (2) Perturb-seq CRISPR KO data from primary melanoma cells under standard mono-culture and co-culture with autologous TILs (53). Processed counts data was downloaded from https://singlecell.broadinstitute.org/single_cell/study/SCP1064/multi-modal-pooled-perturb-cite-seq-screens-in-patient-models-define-novel-mechanisms-of-cancer-immune-evasion. (3) CRISPR activation Perturb-seq screen in K562 cells (51), downloaded from GEO (GSE133344). For each dataset counts were converted to transcript per million (TPM) values as described above and two-sided t-tests were performed to identify differentially expressed genes for each perturbation in each one of the screens, comparing the cells with the perturbation to those with control sgRNAs. MTIL overall expression (OE, **Supplementary Information**) was computed, and a two-sided *t*-test was performed to examine if OE was significantly higher or lower in the cells with the perturbation compared to the control cells (with control sgRNAs). For perturbations that showed a significant effect on the MTIL OE (BH FDR < 0.05, t-test), hypergeometric tests were used to further confirm that the perturbation significantly repress or activates the MTIL genes, having opposite effects on the MTIL-up and MTIL-down gene subsets.

### Transduction and pooled gene knockouts in ovarian cancer cells

#### Plasmid amplification

LentiCas9-Blast was a gift from Feng Zhang (Addgene, Plasmid #52962; RRID:Addgene_52962). Stbl3 bacterial stab was streaked onto a LB broth plate with ampicillin and incubated overnight at 37℃. A single colony was picked and inoculated in 3 mL of LB liquid media supplemented with ampicillin and incubated overnight at 37℃ with 300 rpm agitation. An aliquot of starter culture was then cultured overnight in 10 mL of LB liquid media supplemented with ampicillin. Plasmid extraction was performed using the NucleoBond Xtra Midi EF kit (Takara Bio, 740420.10).

#### Pooled sgRNA Perturb-seq library design

Individual guide sequences were selected from the Human CRISPR Knockout Pooled Library (GeCKO v2) (85). The pooled sgRNA library was purchased from GenScript in a plasmid format utilizing the pLentiGuide-Puro vector. In total, it includes 232 sgRNAs targeting 74 genes, with three guides per gene including ten non-targeting controls (**Table S7**).

#### Lentivirus production

To obtain lentiviral stocks of lentiCas9-Blast and the pooled lentiviral library, LentiCas9-Blast and the custom sgRNA lentiviral library were transfected into Lenti-X 293T cells (Takara, 632180) respectively. Lenti-X 293T cells were cultured in cOPTI-MEM (opti-MEM, Gibco, 31985088), 1x GlutaMAX (Gibco, 35050061), 1 mM Sodium Pyruvate (Corning, 25-000-Cl), 5% FBS, 1x non-essential amino acid (Corning, 25-025-CI). At ∼90% confluency, cells were incubated with TransIT-Lenti (MirusBio, 6603) transfection mixture at 37℃ with 5% CO2. The transfection mixture included cOPTI-MEM supplemented with 14 µg of the respective transfer plasmid, 10 µg psPAX2 (Addgene, Plasmid #12260), and 4.33 µg pMD2.G (Addgene, Plasmid #12259). After six hours of transfection, the media was replaced with fresh cOPTI-MEM supplemented with 1x ViralBoost (Alstem Bio, VB100) and incubated for an additional 16 hours. The supernatant was harvested 24 and 48 hours post-transduction. Harvested viral supernatants were pooled and concentrated with Lenti-X Concentrator (Takara Bio, 631232) by centrifugation at 1,500 x g for 45 minutes. Viral pellets were resuspended in media at a volume 100x smaller than the original volume and stored at −80℃ until retrieved for experiments.

#### Cas9 and sgRNA lentiviral transductions in TYK-nu cells

To obtain stable Cas9 expression in TYK-nu cell line, 100,000 wildtype TYK-nu cells were seeded in a 24-well plate (Corning, 3526) and incubated overnight. Cells were transduced with the lentiCas9-Blast lentivirus at an MOI of 0.2 with 8 _µ_g/mL of polybrene (MilliporeSigma, TR-1003) and incubated overnight in 37℃ with 5% CO_2_. Transduced TYK-nu cells were then washed with DPBS, and post-transduction selection was conducted over ten days with 10 µg/mL of Blasticidin (Invivogen, ant-bl-05) supplementation in TYK-nu media. Successful transduction of Cas9 was validated via western blot (**Figure S6a,c**) and flow cytometry analyses (**Figure S6b,d**). The same process was performed for the custom sgRNA lentiviral library transduction. Post-transduction selection was conducted over five days with 0.5 µg/mL of puromycin (Invivogen, ant-pr-1) at an MOI of 0.15.

To validate knockout efficiency in the TYK-nu Cas9 cell line, TYK-nu Cas9 cells were transduced with pMCB306 (a generous gift from the Bassik Lab), which contains a Puromycin-T2A-EGFP with EF-1 alpha promoter and an EGFP-targeting sgRNA driven by a mU6 promoter. Following cell transduction with the pMCB306 plasmid, loss of GFP fluorescence indicates functional Cas9 activity, as cleavage of GFP by Cas9 results in loss of fluorescence whereas intact GFP retains fluorescence. TYK-nu Cas9 cells were transduced at an MOI of 0.15 with pMCB306 virus and 8 µg/mL of polybrene and incubated overnight at 37℃ with 5% CO_2_. Transduced TYK-nu Cas9 cells were washed with DPBS, and selection was conducted over five days with 0.5 µg/mL of puromycin (Invivogen, ant-pr-1) supplemented in TYK-nu media. Successful transduction was validated via flow cytometry and Western Blot analyses.

#### Flow cytometry analysis of TYK-nu cells

Flow cytometry analysis was conducted to sort and analyze TYK-nu Cas9 B2M knockout cell line and a TYK-nu Cas9 GFP transduced cell line. For flow cytometry of the TYK-nu Cas9 B2M KO cell line, cells were washed in 1X PBS and stained with Alexa Fluor 700 anti-human B2M antibody (BioLegend, 395708; 1:20 dilution) for 20 minutes. Additional cells were set aside to use as unstained controls and to adjust gating. The cells were washed twice in PBS with 1.5% FBS after staining and were filtered through a 35 µm cell strainer prior to analysis. Flow cytometry was performed on the LSRII instrument in the Stanford Shared FACS Facility. For flow cytometry of the TYK-nu Cas9 GFP cell line, cells were resuspended in 1X PBS, washed twice in PBS with 1.5% FBS, and filtered through a 35 µm cell strainer prior to analysis on a Sony Biotechnology SH800S Cell Sorter at the Chan Zuckerberg Biohub Stanford location. All plots were generated with FlowJo Version 10.8.1.

### Co-culture system for high content CRISPR screens

Co-cultures of TYK-nu and NY-ESO-1 TCR^+^ CD8 T cells. Procedures for generating TYK-nu Cas9 cells with stable expression of the NY-ESO-1 antigen and for primary human CD8 T cell isolation and TCR-engineering are described in **Supplementary Information**. Following isolation and editing of CD8 T cells and TYK-nu cells, co-culture experiments were performed as follows. 4,800 NY-ESO-1 Cas9 TYK-nu cells were seeded into a clear-bottom, black-walled 96-well plate and incubated in 100 µl of T cell medium per well overnight. The next day, CD8 T cell activating Dynabeads were magnetically removed from the NY-ESO-1 TCR^+^ CD8 T cells. The activated NY-ESO-1 TCR^+^ T cells were added at varying effector-to-target ratios (E:T) to a total volume of 100 µL per well. The cells were co-cultured for 24 to 72 hours. TYK-nu cell viability and IFNγ levels in the co-culture were measured to validate the cytotoxicity of the edited CD8 T cells. At the end of each co-culture period, supernatants were collected from the 96 well plate, spun down at 400 x g for five minutes, and stored at −20°C in single use aliquots for subsequent ELISA assays (**Figure S5h**). Each well was washed twice with 200 µl of DPBS to remove the T cells. Following the manufacturer’s protocol, PrestoBlue cell viability dye diluted in T cell medium mixtures were added to each well and incubated for 30 minutes prior to fluorescence plate reader reading (Tecan Infinite M1000). The supernatants collected were diluted 1:1000 prior to the IFNγ ELISA assay (BioLegend, 430104). The same co-culture procedure was performed in parallel with wildtype CD8 T cells from the same donor as a control.

#### Co-cultures of TYK-nu and NK-92 cells

TYK-nu ovarian cancer cells (JRCB Cell Bank, JCRB0234.0) were cultured in EMEM (ATCC, 30-2003) with 10% heat-inactivated FBS (Life Technologies, A3840102). NK-92 cells (ATCC, CRL-2407) were cultured in RPMI 1640 Medium, GlutaMAX Supplement, HEPES medium (Gibco, 72400-047) with 10% heat-inactivated FBS (Life Technologies, A3840102), 200 U/mL recombinant human IL-2 (PeproTech, 200-02), and 1% penicillin-streptomycin 100X solution (Cytiva, SV30010). NK-92 cell line cytotoxicity was validated using PrestoBlue cell viability dye (Thermo Scientific, A13261) following the manufacturer’s protocol. TYK-nu cells were seeded in a clear bottom, black walled 96 well plate (Greiner, 655090) and incubated overnight. NK-92 cells were added in varying effector to target ratios (E:T) and the cells were incubated for 24 to 72 hours (**Figure S6e**). To validate the specificity of NK cell cytotoxicity in the co-culture experiments, a TYK-nu B2M KO cell line was generated. B2M is a subunit of the major histocompatibility complex 1 (MHC-1), and its knockout increases cell susceptibility to NK-mediated cytotoxicity by disabling MHC-1 interactions with inhibitory KIRs and NKG2A (86). TYK-nu Cas9 cell lines were transduced with B2M sgRNA lentivirus at an MOI of 0.15. Successful transduction was validated via flow cytometry and western blot analyses (**Figure S6c-d**). All cell lines were routinely tested for mycoplasma using the Promokine PCR Mycoplasma Test Kit I/C (PromoKine, PK-CA91-1024).

#### CRISPR screen in cancer CD8 T cell co-cultures

TYK-nu Cas9 NY-ESO-1 cells were transduced with the sgRNA library at an MOI of 0.15. NY-ESO-1 expressing TYK-nu Cas9 cells were selected with 0.5 µg/mL of puromycin over a period of five days. 1,875,000 NY-ESO-1 expressing TYK-Nu Cas9 library cells were seeded in a 75 mm dish and allowed to adhere overnight to achieve 8,000X coverage. NY-ESO-1 TCR^+^ CD8 T cells were added at 5-to-1 effector-to-target cell ratio (9,325,000 cells). TYK-nu Cas9 NY-ESO-1 cells were grown in: (1) monoculture, (2) co-culture with wild type (WT) CD8^+^ T cells, and (3) co-culture with NY-ESO-1 TCR^+^ CD8 T cells. In all three conditions TYK-nu cells were incubated for 72 hours, either in monoculture or co-culture, before being washed twice in 1X DPBS to remove the CD8 T cells. TYK-nu cells were snap frozen and stored at −80℃ prior to genomic DNA extraction and sgRNA amplification. As a second selection, two days after recovery, cells were grown under the same conditions again for 72 hours and then allowed to recover again before collection and sequencing. All samples were sequenced on a MiSeq Micro V2 in a single-end run at the Chan Zuckerberg Biohub Stanford location.

### CRISPR and Perturb-seq screens in cancer NK co-culture models

TYK-nu Cas9 cells were transduced with the HGSC sgRNA library at MOI of 0.15. TYK-nu Cas9 cells were puromycin selected at 0.5 µg/mL for five days and allowed to recover to confluency prior to downstream experiments.

The first screen was performed for sgRNA and Perturb-seq readouts. TYK-nu Cas9 library cells were seeded in a 75 mm dish (Corning, 353136) and allowed to adhere overnight. NK-92 cells were added at 1-to-1 and 1-to-2.5 effector-to-target (E:T) cell ratios. Perturb-Seq readouts (53,87) were obtained from TYK-nu Cas9 cells grown for 48 hours in monoculture and co-culture with NK-92 cells. After the completed growth timeline, TYK-nu Cas9 library cells were washed twice with 10 mL 1X DPBS to remove the suspended NK-92 cells. Two replicates from each condition were put into a single cell suspension according to the 10X Single Cell Suspensions from Cultured Cell Lines for Single Cell RNA Sequencing protocol (10X Genomics, CG00054 Rev B). The libraries were prepared by the Stanford Genomics Service Center according to the Chromium Next GEM Single Cell 5’ Reagent Kits v2 (Dual Index) with Feature Barcode technology for CRISPR Screening protocol (10X Genomics, CG000510 Rev B). Equimolar amounts of indexed libraries were pooled together and sequenced on a NextSeq2000 P3 in a paired-end run at the Chan Zuckerberg Biohub Stanford location.

A subset of replicated cells was allowed to recover for an additional day until confluency prior to being snap frozen and stored at −80℃. Genomic DNA of the snap frozen cells was extracted using the Quick-DNA Midiprep Plus Kit (Zymo Research, D4075). sgRNA amplification was performed following a previously published protocol(49). Equimolar amounts of indexed libraries were pooled together and sequenced on a MiSeq Nano V2 in a single-end run at the Stanford Genomics Service Center.

A second screen was performed for sgRNA sequencing. TYK-nu Cas9 library cells were seeded in a 6 well dish (Cole-Parmer, 0192770) and were allowed to adhere overnight. NK-92 cells were added at a 2.5-to-1, 5-to-1, and 7.5-to-1 effector-to-target ratios for 48 hours. TYK-nu cells were allowed to recover for three days prior to being snap frozen and prepared for genomic DNA extraction as described above. Each experimental condition was performed in triplicates with > 1000x cells per sgRNA, resulting in 6 and 12 sequencing samples from the first and second screen.

#### CRISPR screen and Perturb-seq data analyses

Raw fastq files were processed using the cellranger pipeline (10x Genomics Cell Ranger 7.1.0). Counts were converted to transcript per million (TPM) values. For each condition (monoculture, 1:1 co-culture, and 2.5:1 co-culture) data was analyzed to remove non-malignant cells. Seurat R package was used for KNN clustering, resulting in a distinct NK cluster in the co-culture conditions, with expression of *CD3E* and *NCAM1*. This cluster was removed and only cancer cells with a detection of a single sgRNA were retained for downstream analyses. For each of the three conditions, DEGs were identified for each perturbation using a two-sided t-test comparing the cells with the perturbation to those with NTCs. Fisher test was used to combine the three p-values. Hypergeometric tests were performed to examine if the up or down regulated genes identified for each perturbation were enriched with M_TIL_-up or M_TIL_-down genes, or vice versa, and the combined p-value (fisher test) was reported as the final summary statistics.

MAGeCK algorithm was used to compute differential fitness effects on the cancer cells under the monoculture and co-culture conditions, either with the different types of CD8^+^ T cells or with the NK cells. Each experimental condition was performed in triplicates. First, the sgRNA counts of the different samples were median normalized to adjust for the effect of library sizes and read count distributions. Second, the variance of read counts was estimated by sharing information across the different sgRNAs, allowing to fit a negative binomial (NB) model to test whether sgRNA abundance differs significantly between treatments (i.e., co-culture) and controls (i.e., monoculture or co-culture with non-specific T cells). Third, sgRNAs were ranked based on p-values calculated from the NB model, and an α robust ranking aggregation (α-RRA) algorithm was used to identify positively or negatively selected genes. The pairwise tests were performed considering each of the co-cultures (CD8 T cell or NK cell) compared to the monoculture and the two co-cultures (with specific and non-specific T cells) to compute a combined Fisher statistics, one for T cell and another for NK cell sensitizing and desensitizing hits.

## DATA AVAILABILITY

All the data collected in this study, including spatial transcriptomics data, single-cell Perturb-seq data, targeted genomics, deidentified clinical meta-data, and processed tissue images will be deposited and made publicly accessible through Gene Expression Omnibus (GEO), Zenodo, and CELLxGENE. Processed data in the form of standardized RObjects will be available via Zenodo. Upon publication raw and processed spatial transcriptomics data will be available on CELLxGENE for download in .h5ad format and interactive exploration. Processed gene expression matrices with cell type annotations from 6 scRNA-seq studies with HGSC tumor samples were downloaded from publicly available repositories specified in their respective publications (23,27–32). Specifically, preprocessed gene expression and metadata matrixes of HGSC scRNA-seq data were downloaded from Synapse (syn33521743 (23)), GEO (GSE118828 (27), GSE173682 (29), GSE147082 (28), GSE154600 (31), GSE146026 (80)), and https://lambrechtslab.sites.vib.be/en/data-access (30,32). An additional external validation dataset (44) hosted on the European Genome-Phenome Archive (EGAD00001006973, EGAD00001006974) was made available for this study through a Data Access Agreement with Genentech, Inc..

## CODE AVAILABILITY

All data processing and analysis code will be available via GitHub once the paper is published. The study GitHub repository includes documented code for data processing and code required to reproduce the main figures and supplementary tables of the study.

## ACKNOWLEDGEMENTS

We thank Reece Villarin Akana, Georgia Schmitt, Grace Allard, and Kaitlyn Spees for help with setting up assays and models related to this work; Lettie McGuire for help with artwork; Kai Wucherpfennig and Nathan Mathewson for generously providing NY-ESO-1 constructs; Theodore Roth for discussions on primary T cell culture; Stanford Genomics Service Center (SFGF), Norma Neff and the rest of the Stanford Chan Zuckerberg Biohub Center team for ongoing support. L.J. is a Chan Zuckerberg Biohub Investigator and an Allen Distinguished Investigator. L.J. holds a Career Award at the Scientific Interface from the Burroughs Wellcome Fund and a Liz Tilberis Early Career Award from the Ovarian Cancer Research Alliance (OCRA). This study was supported by the OCRA The Liz Tilberis Early Career Award Grant 889076. This work was support by Under One Umbrella, Stanford Women’s Cancer Center, Stanford Cancer Institute, an NCI-designated Comprehensive Cancer Center, and supported in part by funds from the Departments of Pathology and Genetics at Stanford University. Processed spatial and scRNA-seq Perturb-seq data will be available on the Gene Expression Omnibus (GEO) and Zenodo. Figure 1a, 7a and Figure S6a include elements created with BioRender.com (Licensing Agreement Number: AB25KWR1LU).

## Author Contributions

C.Y.Y., K.A., O.L., B.E.H, and L.J., designed the study. C.Y.Y., K.A., O.L., B.E.H, and L.J. conducted the research and interpreted the results. K.A., O.L., and S.K., preformed the experiments with L.J. supervision. A.W., B.L., X.Z., L.H., and B.E.H. collected, annotated, and provided tissue samples for profiling. C.Y.Y., and L.J. performed the computational and statistical analyses. R.V. supported experimental design. M.C.B. supported experimental design and manuscript writing. M.P.S. supported data collections. S.K.P. supported the computational and statistical analyses. C.Y.Y., K.A., O.L., B.E.H, and L.J., wrote the manuscript. B.E.H. and L.J. supervised the study. B.E.H. and L.J. obtained funding. All authors reviewed and approved the manuscript.

---

**Figure S1.**
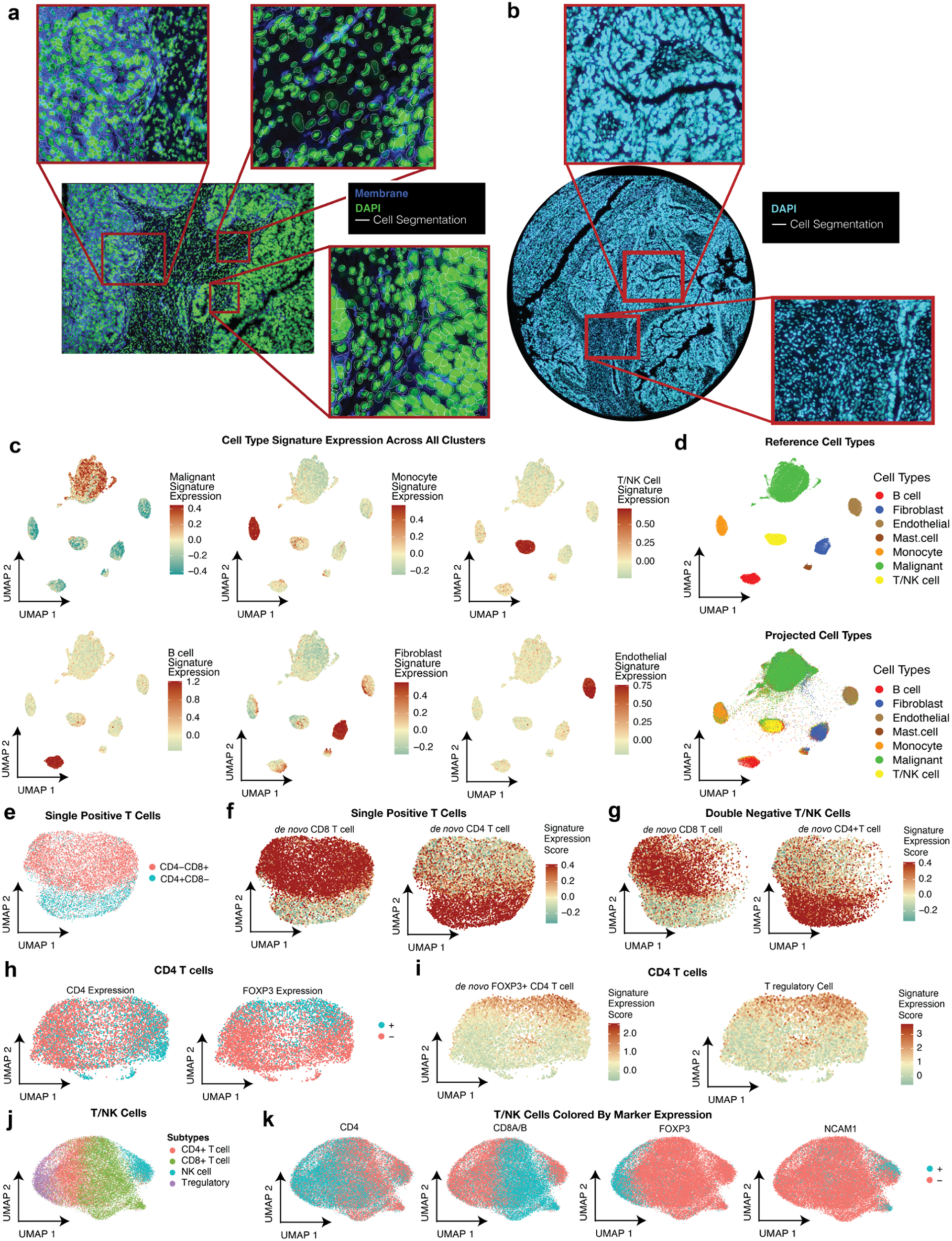
Cell segmentation and cell type annotations of spatial transcriptomics. **(a)** Representative whole-cell segmentation performed for the discovery dataset. Input data includes DAPI immunofluorescent (IF) stain (green) and cell membrane stain (blue). Cell boundaries represented as white contours. (**b)** Representative nuclear segmentation performed for validation dataset 1. Input data includes DAPI IF stain. Cell boundaries represented as white contours. **(c)** UMAP of cell transcriptomes, cells colored by overall expression of cell type signatures (**Supplementary Table 3, Methods**) corresponding to the following cell types: malignant, monocyte, T/NK cell, B cell, fibroblast, and endothelial cells. (**d)** Reference UMAP embedding fit of single cell transcriptomes from the discovery dataset, shown for a subsample with high confidence cell type annotations (top) and all cells projected onto the reference embedding (bottom), colored by cell type annotations. (**e-f)** UMAP embedding of single positive T cell transcriptomes in the discovery dataset, cells colored by (**e)** CD8 and CD4 expression, and (**f)** expression of *de novo* CD8 (left) and CD4 (right) T cell expression signatures. (**g)** Projection of double negative T/NK cell transcriptomes onto UMAP embedding in (e), with cells colored by overall expression of the *de novo* CD8 (left) and CD4 (right) T cell gene signatures (**Supplementary Table 3**). (**h)** UMAP embedding of CD4 T cell transcriptomes, cells colored by CD4 expression (left) and *FOXP3* expression (right). (**i)** UMAP as in (h), with cells colored with *de novo* FOXP3^+^CD4 T cell gene signature expression score (left) and cells colored with the regulatory T cell signature derived from publicly available scRNA-Seq datasets (**Methods**; **Supplementary Table 3**). (**j-k)** UMAP embedding of validation dataset 1 T/NK single cell transcriptomes, cells colored by (**j)** T/NK cell subtype annotations, (**k)** detection of (from left to right): *CD4*, *CD8A/B*, *FOXP3* (regulatory T cell marker), and *NCAMI* (NK cell marker). All signatures used in and generated by these analyses are provided in **Supplementary Table 3.**

**Figure S2.**
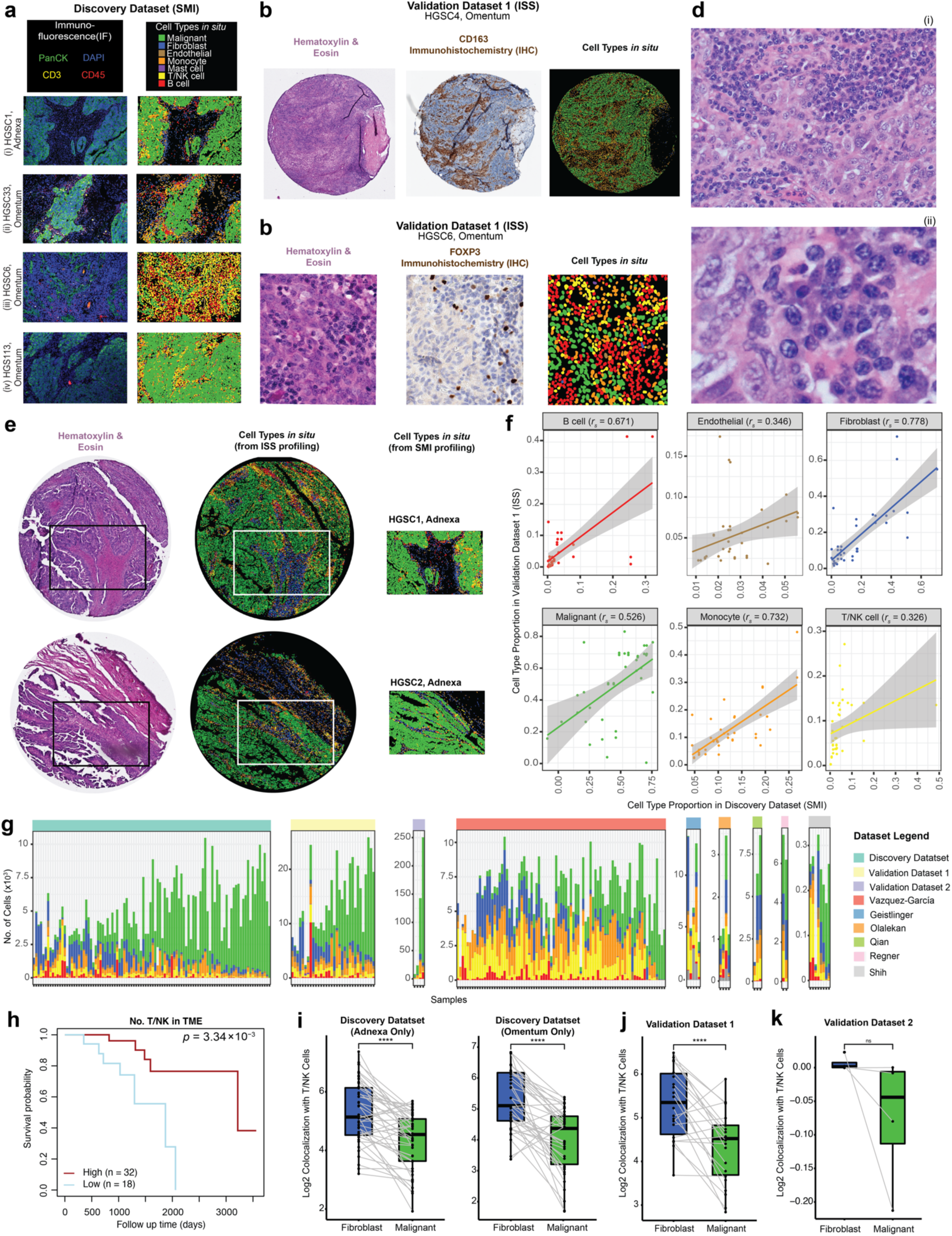
Cross-platform validation and evaluation of cell type annotations, compositions, and tumor architecture. **(a)** Immunofluorescence (left column) of cell type markers paired with cell type annotations plotted *in situ* (right column) for four representative patient samples (rows) in the discovery dataset. (**b-g)** Cell types colored according to cell type legend in (a). (**b)** Hematoxylin & Eosin staining (H&E, left), Immunohistochemistry (IHC) stain for CD163 (middle; monocyte marker) with corresponding cell type annotations *in situ (*right) in a representative tissue FOV in validation dataset 1. (**c)** H&E (left), IHC stain for FOXP3 (middle, Treg marker), and corresponding cell type annotations *in situ (*right) in one representative tissue FOV from validation dataset 1. (**d)** High power H&E stains of HGSC6 omentum tumor tissue resolving morphology of plasma cells identified based on the discovery cohort in this sample as shown in panel a (iii). (**e)** H&E (left), annotated cell types *in situ* from ISS validation dataset 1 (middle) and SMI discovery dataset (right) showing matching data (same patient, same tumor) from two tumors (rows). White box denotes region of tissue profiled by ISS that corresponds to FOV profiled by SMI in the same row. (**f)** Cell type proportion in biological replicates profiled by both SMI (x axis) and by ISS (y axis). Straight lines correspond to the linear regression fit. *r_s_* denotes the Spearman correlation coefficient. (**g)** Stacked barplot show the number of cells (y axes) profiled stratified by cell type (color) and shown for the individual samples (x axes) and datasets (panes, labeled by ST platform name or first author of published scRNA-seq dataset)(27,29–32,72,73). (**h**) Kaplan Meier Survival curves depicting differential survival probability (y axis) as a function of average T/NK abundance in each patient (log rank test *p* = 3.34*10^-3^). **(i-k)** Log_2_ Co-localization Quotient (CLQ, y axes) of T/NK cells with fibroblasts (blue, x axis) and T/NK cells with malignant cells (green, x axis) in (**i)** the discovery dataset, stratified by adnexal samples (left) and omentum samples (right), (**j)** all samples in validation dataset 1 (**k)**, all 4 tissue section samples in validation dataset 2 (all adnexal). Light grey lines connect paired fibroblasts and malignant cells derived from the same tissue section. *\*\*p* < 1*10^-2^, *\*\*\*p* < 1*10^-3^, *\*\*\*\*p* < 1*10^-4^, paired Wilcoxon rank sum test. Boxplots middle line: median; box edges: 25^th^ and 75^th^ percentiles; whiskers: most extreme points that do not exceed ± IQR x 1.5; further outliers are marked individually.

**Figure S3.**
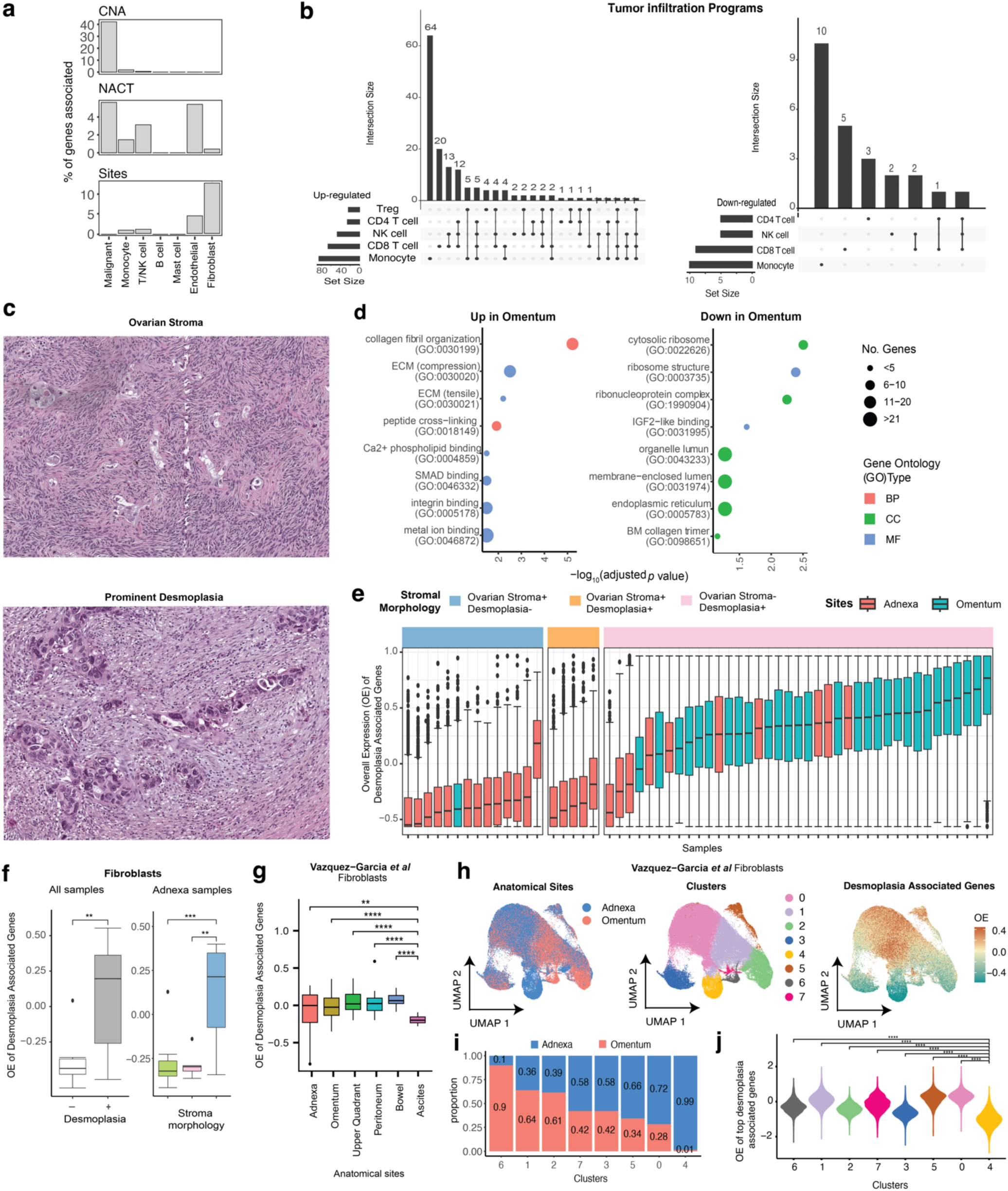
Cellular programs linked to anatomical sites and desmoplasia. **(a)** Percentage of genes (y axis) that are significantly associated with (*p* < 0.05, mixed effects BH FDR) somatic copy number alterations (CNA; top), treatment status (NACT = neoadjuvant chemotherapy; middle), and tumor anatomical site (i.e., adnexa or omentum; bottom) in the discovery dataset. (**b)** Size (horizontal bars) and overlap (vertical bars) between the tumor infiltration programs identified for the five different immune cell subsets, shown for the up-regulated (left) and down-regulated (right) subsets. (**c)** H&E of normal ovarian stroma morphology (left), and desmoplastic stroma morphology (right). (**d**) Gene ontology enrichment analysis (**Methods**) of the top desmoplasia associated genes. Abbreviations: BP = biological process, CC = cellular component, MF = molecular function. (**e-g)** overall expression (OE) of desmoplasia associated genes (**Supplementary Table 5**) per fibroblast (y axis) in (**e)** discovery dataset, stratified by sample, (**f)** discovery dataset, per sample (y axis) as a function of stromal morphology annotations (x axis) across all samples (left) and in adnexal samples only (right). \*\**p* < 1*10^-2^, \*\*\**p* < 1*10^-3^, mixed effects test. **(g)** fibroblasts the Vazquez-Garcia *et al*(72) scRNA-seq dataset, stratified by anatomical site \*\**p* < 1*10^-2^, \*\*\*\**p* < 1*10^-4^. (**h)** UMAP embeddings of adnexal and omentum fibroblasts from the Vazquez-Garcia *et al* (72) scRNA-seq dataset with each cell colored by anatomical site (left), unsupervised shared nearest neighbors clusters (middle), and the OE of desmoplasia associated genes (right) (**i)** proportion of cells (y-axis) from the adnexa vs. omentum in each cluster (x-axis) as defined in (h). (**j)** OE of desmoplasia associated genes (y-axis) in each cluster (x-axis) as defined in (h), \*\*\*\**p* < 1*10^-4^.

**Figure S4.**
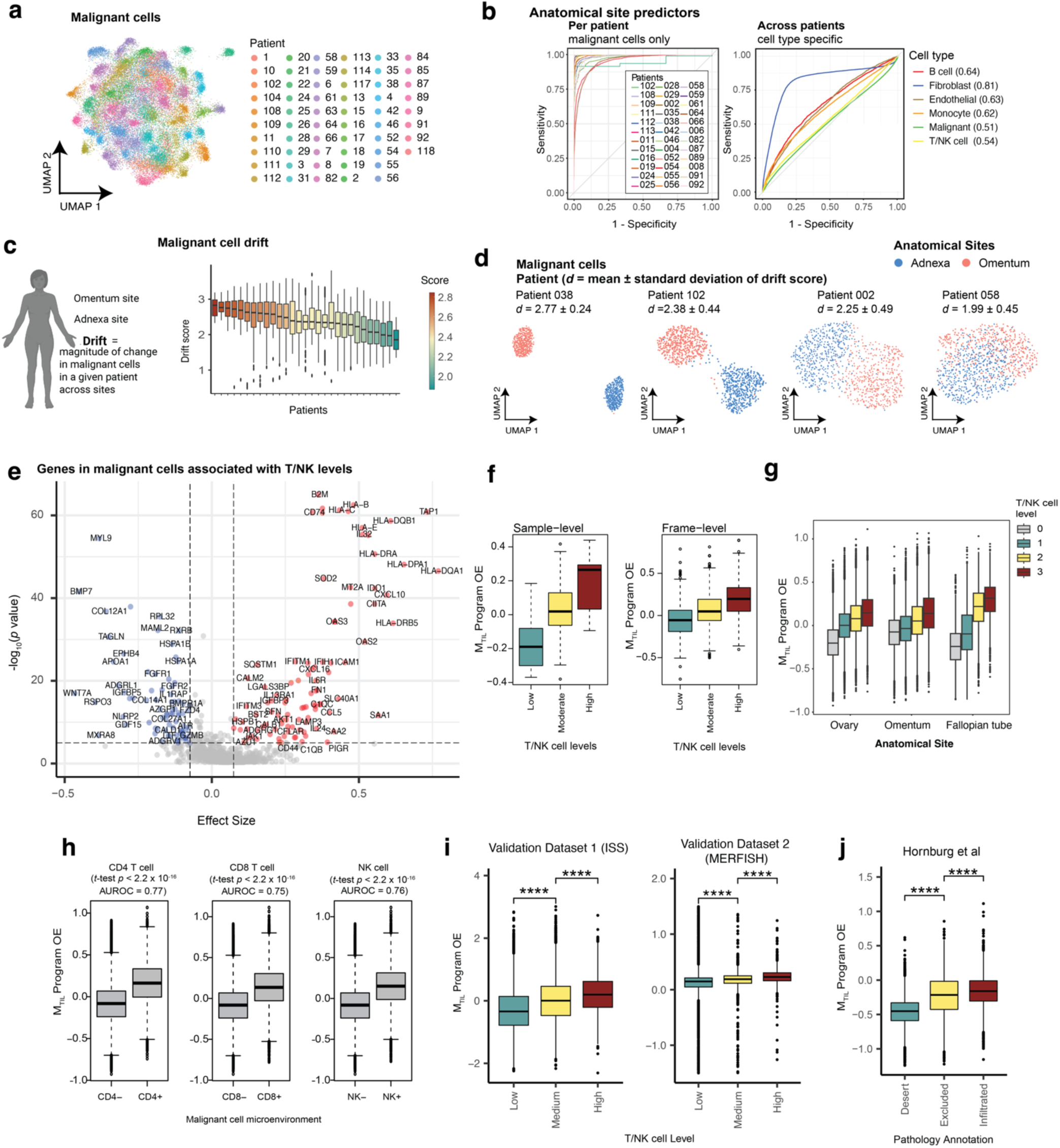
M_TIL_ marks T/NK infiltration across multiple scales and datasets. **(a)** UMAP embedding of malignant cell ST profiles from the discovery dataset, colored by patient. **(b)** Receiver Operating Characteristic (ROC) curve obtained for Random Forest (RF) classifiers trained to predict if a cell was obtained from adnexal or omentum tumors. Area under the receiver operator curve (AUROC) is reported in parenthesis. **Left**: Patient-specific RF classifiers trained to predict the anatomical site of malignant cells. Each classifier was trained per patient and tested on unseen malignant cells from the same patient. **Right**: Cell type specific RF classifiers trained per cell type and tested on unseen patients. (**c)** Variation in malignant cell gene expression “drift” score (y axis, **Supplementary Information**) across patients with paired adnexa and omentum tumor samples. (**d)** UMAP embedding of malignant cell ST profiles from the adnexa (blue) and omentum (pink), depicted for four representative patients. The magnitude of the malignant gene expression drift identified per patient is denoted by *d* (**Supplementary Information**). (**e)** Significance (y axis) and effect size (x axis) of association of malignant gene expression with T/NK levels quantified via mixed effect models in the discovery dataset (**Methods**). (**f-h)** Discovery dataset: M_TIL_ overall expression (OE; y axis) as a function of (**f)** discretized T/NK levels (x axis) across samples (left) and spatial frames (right), (**g)** T/NK levels (color) and anatomical site (x axis), (**h)** presence of T/NK cell subtypes in the spatial frame: CD4 T cells (left), CD8 T cells (middle), and NK cells (right). AUROC: Area Under the Receiver Operating Characteristic Curve. (**i-j)** M_TIL_ OE (y axis) in malignant cells as a function of (**i)** T/NK levels (x axis) in validation data 1 (left) and validation data 2 (middle), and (**j)** sample immune type (x axis) labeled by expert pathologists from Hornburg *et al* scRNA-seq study (44). In (f-i) boxplots: middle line = median; box edges = 25^th^ and 75^th^ percentiles; whiskers = most extreme points that do not exceed ± IQR x 1.5; further outliers are marked individually.

**Figure S5.**
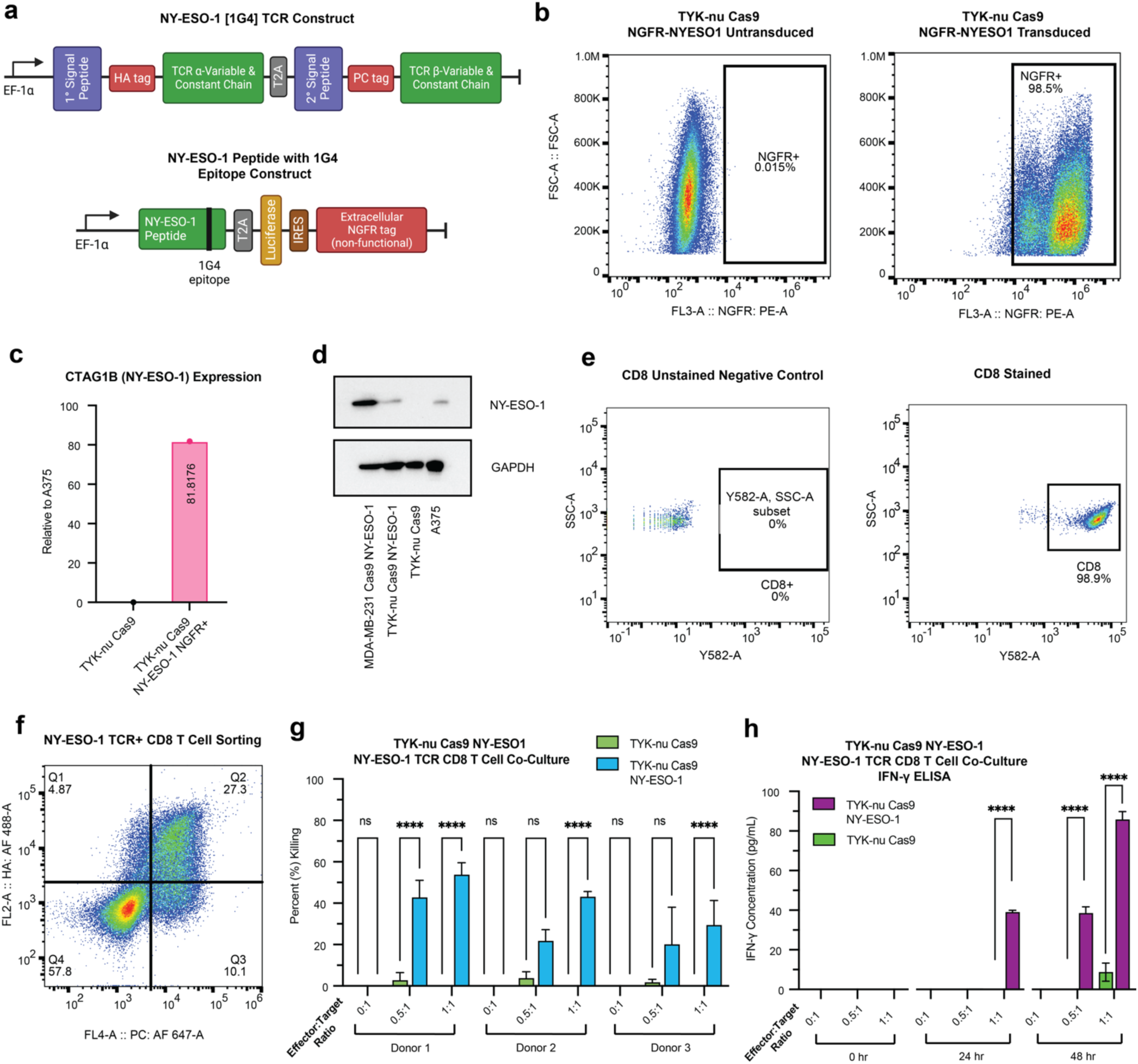
Validation and design of ovarian cancer-CD8^+^ T cell CRISPR screen. (**a)** Top: NY-ESO-1 [1G4] TCR lentiviral construct used to engineer primary human CD8^+^ T cells (88), with α and β-chains tagged by HA and PC tags, respectively. Bottom: NY-ESO-1 peptide with 1G4 epitope lentiviral construct used to edit TYK-nu Cas9 (88) cells to express the 1G4 NY-ESO-1 antigen. A non-functional, extracellular domain of human growth factor receptor (NGFR) was used as a tag to identify and sort NY-ESO-1 expressing cancer cells via flow cytometry. (**b)** Representative flow cytometric analysis gated on the expression of the non-functional NGFR tag to quantify TYK-nu Cas9 cells transduced to express NY-ESO-1 antigen. (**c)** qPCR quantification of *CTAG1B* mRNA expression in NY-ESO-1 transduced TYK-nu Cas9 cell line (TYK-nu^NY-ESO-1+^) relative to A375 melanoma cell line with endogenous *CTAG1B* expression, encoding for NY-ESO1. All data shown represents the mean +/- s.e.m. (**d)** Western blot of NY-ESO-1 expression from NY-ESO-1 transduced MDA-MB-231 Cas9, TYK-nu^cas9,NY-ESO-1+^, TYK-nu Cas9, and A375 whole cell lysates. GAPDH was used as a loading control. (**e)** Representative flow cytometric analysis of CD8^+^ T cells isolated from PBMC of a healthy human adult donor. (**f)** Representative flow cytometric analysis of NY-ESO-1 TCR transduced CD8 T cells. HA (α chain) and PC (β chain) tags double-positive CD8^+^ T cells were sorted via flow cytometry to ensure complete expression of NY-ESO-1 TCR. (**g)** 24-to-72-hour time course T cell co-culture cytotoxicity assay with CD8 T cells from three different donors (x axis). NY-ESO-1 TCR expressing primary CD8 T cells were co-cultured with TYK-nu Cas9 cells or TYK-nu^Cas9,NY-ESO-1+^ cells at variable effector to target cell ratios (E:T). The percentage of killed (PrestoBlue negative) tumor cells was calculated by normalizing to tumor cell monoculture conditions. Co-cultures were performed using 3 replicates per condition and three biological replicates. All data shown represent the mean +/- s.e.m. (**h)** ELISA quantification of IFNγ secreted in the co-culture supernatant (1:1000). Co-culture was conducted in the same manner as described in (g). All data shown represent the mean +/- s.e.m.

**Figure S6.**
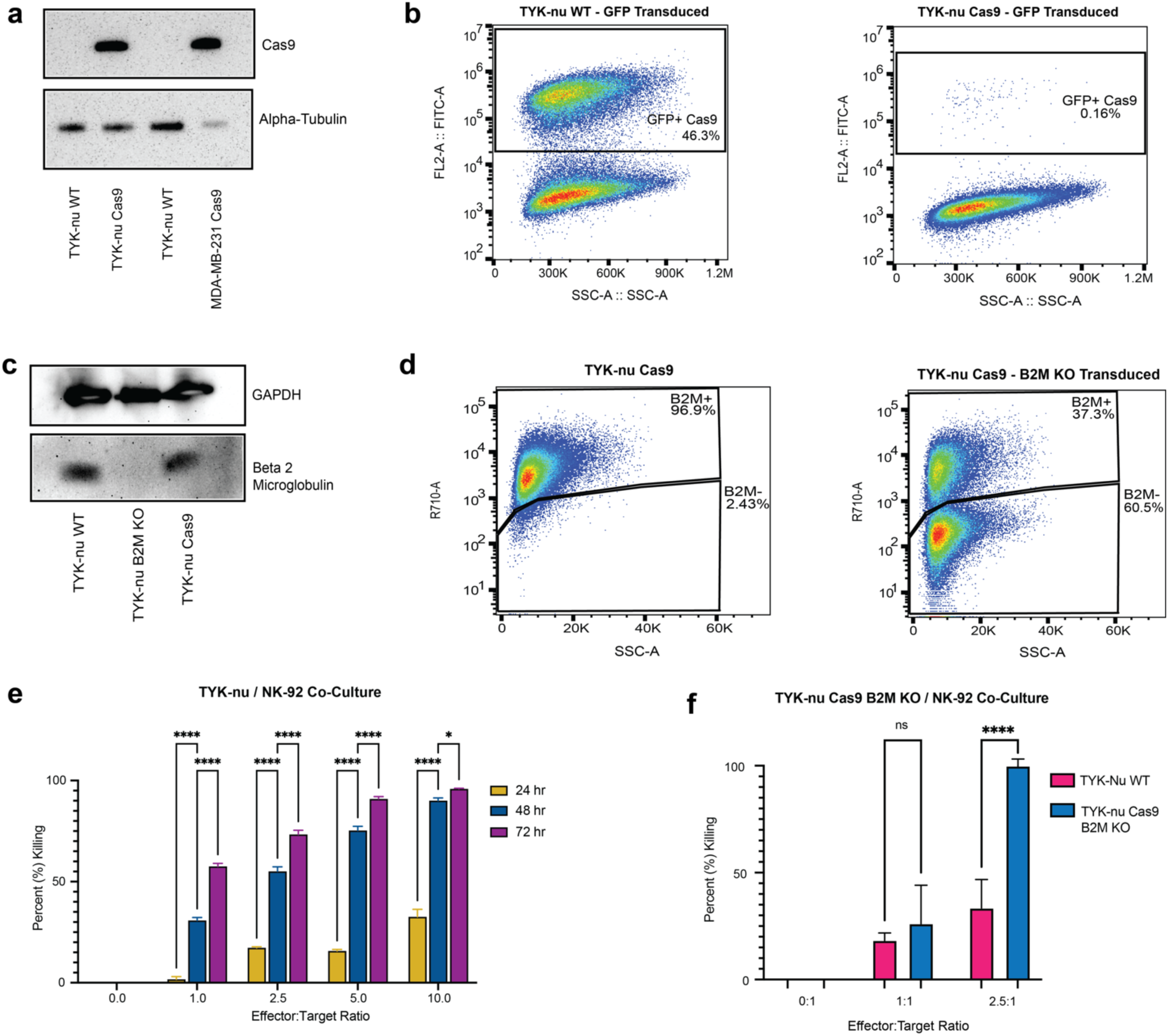
Validation and design of ovarian cancer NK cell co-culture model used in the Perturb-Seq screen. (**a)** Western blot of Cas9 protein from WT and Cas9 transduced whole cell lysates. Alpha tubulin measured as loading control. (**b)** Representative flow cytometric analysis gated on GFP expression to measure Cas9 efficiency using pMCB306 plasmid (**Methods**), comparing GFP levels WT vs. Cas9 TYK-nu cells following pMCB306 transduction. Loss of GFP denotes Cas9 activity. (**c)** Western blot of beta-2-microglobulin (B2M) from whole cell lysates of WT, Cas9, and B2M^KO^ TYK-nu. GAPDH measured as a loading control. (**d)** B2M surface expression by flow cytometry in B2M^wt^ and B2M^KO^ Cas9 TYK-nu cells. (**e)** 24-to-72-hour time course cell viability co-culture with TYK-nu Cas9 and NK-92 cell lines at variable effector to target cell ratios. Percent killing was calculated by normalizing to monoculture conditions. Co-cultures were performed in 4 replicates per condition as shown. (**f)** 48-hour cell viability of B2M^KO^ and B2M^WT^ TYK-nu cell lines in co-culture with NK-92 cells. Percent killing was calculated by normalizing to the monoculture conditions. Co-culture data is represented by the mean +/- s.e.m. and each experiment preformed in four replicates. \*\*\*\**p*< 1*10^-4^, \**p* < 0.05, two-way analysis of variance (ANOVA). All statistical tests were conducted on GraphPad Prism 9.

**Figure S7.**
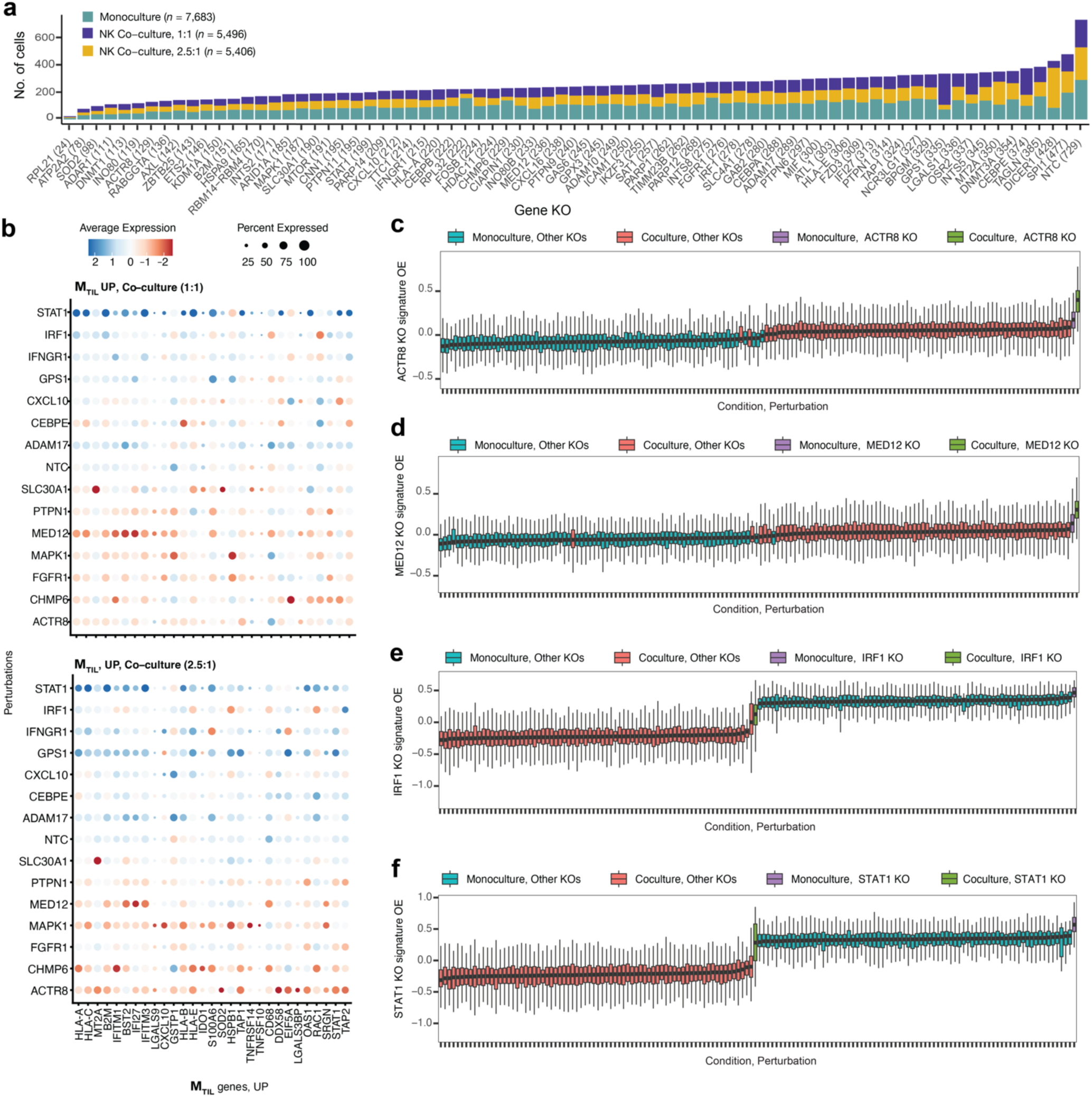
Perturb-seq screen in ovarian cancer identifies immune response regulators. **(a)** Number of cells (y axis) detected with sgRNAs targeting each gene (x axis) in the CRISPR knockout (KO) library. (**b)** Gene expression (color bar) of M_TIL_-up genes (x axis) under different gene knockouts (KO; y axis). Dot color and size represents the average expression and percent of cells expressing the gene, respectively. (**c-f)** Gene KOs mimic (c-d) and repress (e-f) transcriptional response to NK cells: KO gene signature overall expression in each condition and gene KO combination (x axis), shown for *ACTR8* (c) *MED12* (d), *IRF1* (e), and *STAT1* (f).

## SUPPLEMENTAL INFORMATION

**Table S1. Specifications of datasets collected and/or analyzed in this study.**

**Table S2. Metadata for each tissue section profiled by spatial transcriptomics.**

**Table S3. Cell Type Signature Genes.** Includes both gene signature derived from scRNA-seq and CellTypist Immune Encyclopedia (a) and from the HGSC spatial transcriptomics data collected here (b).

**Table S4. Immune tumor infiltration signatures derived from discovery dataset**, shown for different immune cell subsets (a) and for the CD8 T cell infiltration program expanded to whole-transcriptome using scRNA-seq data of CD8 T cells (b).

**Table S5. Fibroblast desmoplasia associated genes** (a) and their Gene Ontology enrichment analysis (b).

**Table S6. MTIL genes** (a) and its Gene Ontology enrichment analysis (b).

**Table S7. High-content CRISPR screens sgRNA library.**

